# Targeting the sinusoidal-to-capillarized transition to normalize vessels in breast cancer liver metastasis

**DOI:** 10.64898/2026.10.28.762442

**Authors:** Han-Ying Huang, Chuang Zhao, Xin-Nan Zheng, Jing-Min Zeng, Yan-Xia Shi, Lin Tian

**Affiliations:** State Key Laboratory of Oncology in South China, Guangdong Provincial Clinical Research Center for Cancer, Sun Yat-sen University Cancer Center, Guangzhou 510060, P. R. China

## Abstract

Liver metastases exhibit marked resistance to anti-VEGF anti-angiogenic therapies, yet the underlying mechanisms remain poorly understood. By combining single-cell RNA sequencing of human breast cancer liver metastases with a liver sinusoidal endothelial cell (LSEC) lineage-tracing mouse model, we show that liver metastasis-associated vessels originate from Lyve1+ sinusoidal endothelial cells but undergo a sinusoidal-to-capillarized transition as they grow into the metastatic nodule, characterized by enhanced inflammatory signaling, loss of sinusoidal markers, and acquisition of capillarized markers. We functionally perturbed each of these abnormalities individually. Unexpectedly, ectopic Lyve1 overexpression paradoxically enhanced hyaluronan and other extracellular matrix deposition in the tumor microenvironment, suggesting a compensatory mechanism. Similarly, targeting TNF signaling, a hallmark inflammatory pathway in liver metastasis, dampened the inflammatory state of metastasis-associated vessels but promoted metastatic outgrowth, indicating that TNF blockade is not a viable strategy for metastasis treatment. Strikingly, inhibition of the capillary tip cell marker Esm1 improved vessel perfusion and enhanced chemotherapeutic drug delivery, two defining features of vascular normalization. Pan-cancer analysis further revealed that Esm1 expression is restricted to tip cells and correlates with poor patient prognosis, positioning Esm1 as a candidate target for promoting vascular normalization. Collectively, our findings reveal the vascular plasticity of liver metastasis and, although not all abnormal features are actionable, we propose targeting Esm1 to inhibit sprouting angiogenesis and to normalize dysfunctional liver metastasis vessels, offering a promising therapeutic avenue for liver malignancies.

## Introduction

Abnormal vasculature is a well-established hallmark of cancer, characterized by aberrant structure and impaired perfusion^1–3^. Anti-angiogenic therapy, which targets the sprouting of new vessels, was originally conceived to starve tumors by disrupting their nutrient supply. Intriguingly, its clinically relevant benefit also derives from pruning non-functional vessels and restoring vascular function, a process termed vessel normalization (VN)^4,5^, thereby enhancing perfusion and drug delivery. To date, VN has been pursued almost exclusively through blockade of the VEGF-VEGFR pathway^6,7^. However, the efficacy of anti-VEGF agents is nevertheless limited in liver metastases, where cancer cells largely co-opt the pre-existing sinusoidal vasculature instead of inducing angiogenesis, a process known as vessel co-option^8,9^. Vessel co-option has been documented in both hepatocellular carcinoma (HCC) and liver metastases (LvMet) of breast and colorectal cancer^10^. Defining the cellular and molecular features of this VEGF-independent tumor vasculature is therefore critical for developing effective vascular-targeting strategies for liver malignancies.

Unlike the vasculature of most tissues, where vessels are ensheathed by perivascular cells and rest on a continuous basement membrane, the hepatic sinusoids are lined by liver sinusoidal endothelial cells (LSECs), a highly specialized endothelial subtype characterized by fenestrae, high permeability, and the absence of a basement membrane. Positioned along the porto-central axis between the portal triads and central veins, LSECs maintain the distinctive exchange function of the liver and exhibit zonation-dependent transcriptional identities^11^. In liver malignancies, however, tumor-associated vessels progressively lose these sinusoidal features and acquire a capillary-like morphology, resembling tumor endothelial cells (TECs) in non-hepatic tumors^12^. Notably, the vascular growth pattern in liver tumors correlates with patient survival, with vessel co-option associated with the poorest clinical outcomes^8,13,14^. Yet whether and how pre-existing LSECs are hijacked and subsequently transform into capillary-like TECs in vivo remains unknown, owing to the lack of definitive lineage-tracing evidence.

Tumors, recognized as “wounds that do not heal,” represent a highly inflammatory environment that has been shown to be either beneficial or detrimental to cancer cells^15^. Beyond their role as passive conduits, tumor vessels function as active signaling platforms that participate in inflammatory pathways^16^. This endothelial activation is particularly relevant in the liver, where LSECs act as immunological gatekeepers that continuously sample blood-borne antigens and modulate leukocyte trafficking, thereby maintaining hepatic immune homeostasis^17^. Inflammatory cytokines such as TNF activate endothelial cells through the NF-κB pathway, inducing adhesion molecules and chemokines that orchestrate leukocyte recruitment and vascular inflammation^18^. Whether blockade of inflammatory signaling could normalize tumor vessels for therapeutic benefit warrants further exploration.

In chronic liver disease, inflammation is tightly coupled to extracellular matrix (ECM) remodeling^19,20^. Inflammatory cytokines and chemokines released during liver injury activate hepatic stellate cells into myofibroblast-like cells that deposit collagen, hyaluronan, and other matrix components, culminating in fibrosis^21,22^. Concurrently, LSECs undergo capillarization, a phenotypic switch characterized by defenestration, basement membrane formation, loss of sinusoidal markers, and acquisition of capillary markers such as laminin and collagen^20^. Notably, these features are mechanistically intertwined. For example, sinusoidal markers such as Lyve1 and Stab2 are scavenger receptors for hyaluronan, and their downregulation results in the loss of anti-fibrotic and matrix-clearing functions, forming a positive feedback loop that further augments hyaluronan accumulation^23^. Given that liver metastases arise in this inflammation- and matrix-rich hepatic context, we hypothesized that a similar inflammation-ECM-capillarization axis may drive vascular dysfunction of liver metastases. Elucidating how these features are causally related in the context of liver metastasis may therefore reveal potential synergistic therapeutic strategies.

In this study, we integrated scRNA-seq data of human liver malignancy samples from multiple resources and filtered high-quality TEC transcriptomes, uncovering a profound phenotypic shift from a sinusoidal to a capillarized state. To confirm this phenotypic plasticity, we employed an autochthonous mouse liver cancer model and an experimental breast cancer liver metastasis model, demonstrating that TECs in both HCC and liver metastases (LvMet) originate from LSECs. During their ingrowth into the metastatic nodule, co-opted vessels progressively lose sinusoidal markers while acquiring inflammatory and ECM-related characteristics. Mechanistically, we tested multiple strategies aimed at normalizing TECs. Restoration of the sinusoidal marker Lyve1 failed to reverse the sinusoidal-to-capillarized transition, and blockade of inflammatory TNFR signaling promoted angiogenesis and potentiated liver metastasis. In contrast, inhibition of Esm1, an ECM-associated tip cell gene, induced VN and improved vessel function. Collectively, our findings elucidate the abnormal vascular features of liver malignancies and identify Esm1 as a prognostic marker for non-angiogenic tumors and a target to promote VN, offering new synergistic opportunities for the treatment of liver tumors.

## Results

### Sinusoidal-to-capillarized transition of tumor vessels in liver malignancies

To dissect the heterogeneity and dysfunction of tumor endothelial cells (TECs), we integrated 501,041 cells from three independent studies^24–26^ (**Fig. 1A**). After selecting endothelial cells based on *VWF* and *PECAM1* (CD31) expression, we obtained 42,910 high-quality TECs from 10 normal liver, 21 hepatocellular carcinoma (HCC), 28 primary breast cancer, and 17 breast cancer liver metastasis samples (**Fig. 1B**). Unbiased re-clustering divided TECs into six clusters defined by endothelial cell-specific signatures^27^. Strikingly, TECs from liver malignancies comprised a mixture of capillarized endothelial cells (CapECs) and liver sinusoidal endothelial cell (LSEC)-like cells, with CapEC proportions exceeding those in normal liver but remaining below those in primary breast cancer, revealing the coexistence of co-opted sinusoidal and angiogenic capillary programs within liver malignancies (**Fig. 1C**). Pseudotime analysis further revealed a sharp loss of sinusoidal markers (*LYVE1*, *STAB2*) accompanied by a gradual acquisition of capillarized markers (*ITGA6*, *LAMB1*) along the trajectory (**Fig. 1D**). This sinusoidal-to-capillarized transition was also observed in colorectal cancer liver metastases, as revealed by both spatial and bulk transcriptomic analyses of patient samples, indicating that it is a general feature of liver malignancies (Supplementary Fig. S1A,B).

**Figure 1.**
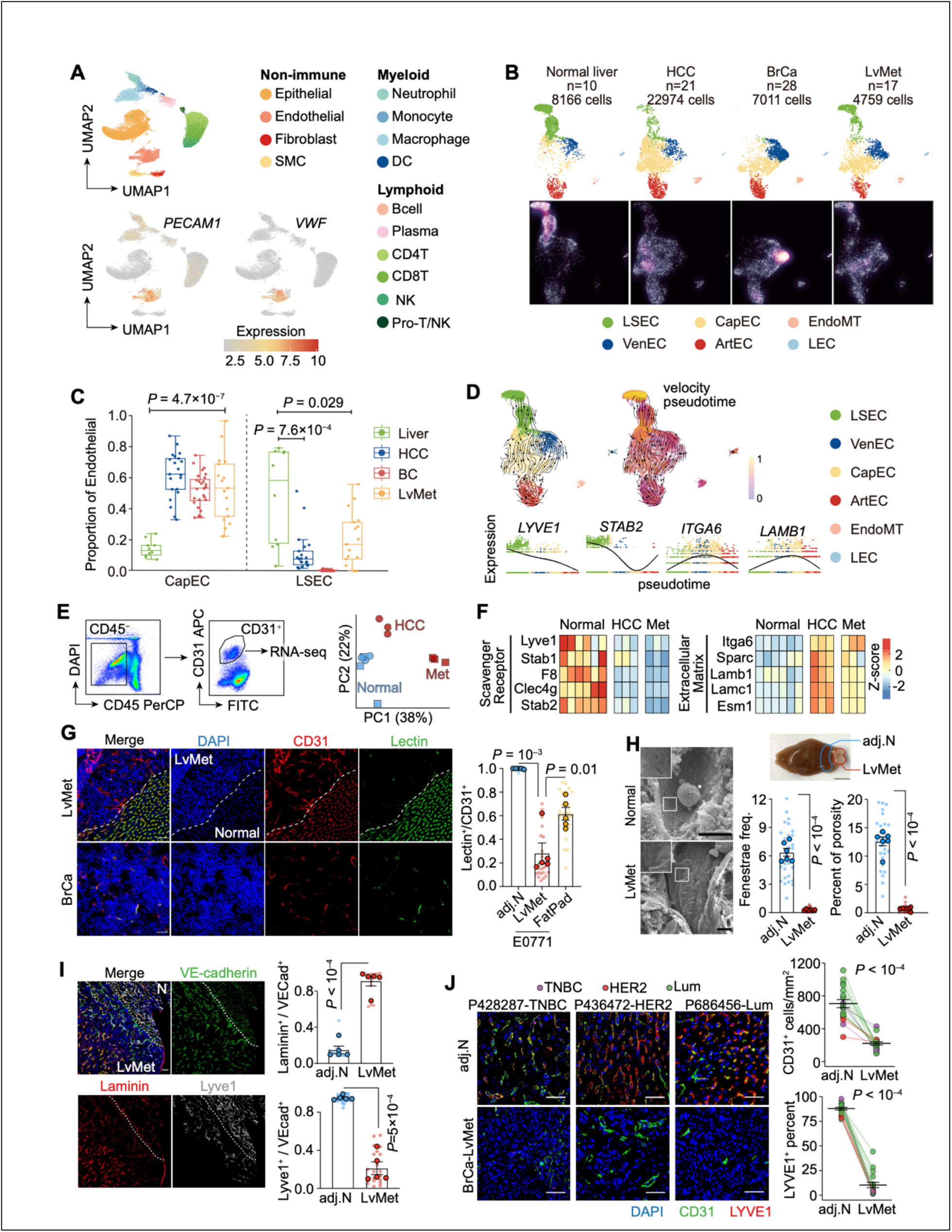
Loss of sinusoidal and gain of capillarized features in vessels of breast cancer liver metastasis. **A,B.** Integration of human scRNA-seq data across tissues. PECAM1 and VWF were used to identify endothelial cell (EC) clusters (**A**), which were further subclustered for endothelial subset annotation (**B**) (HCC, hepatocellular carcinoma; BrCa, primary breast cancer; LvMet, liver metastasis of breast cancer). **C.** Comparison of proportion of sinusoidal EC and capillarized EC across four different tissues. **D.** RNA velocity and pseudotime analysis showing changes in expression of selected sinusoidal genes (LYVE1, STAB2) and capillarized genes (ITGA6, LAMB1) as the endothelial cells transdifferentiate from LSEC to VenEC/CapEC and finally to ArtEC. **E.** Flow cytometry gating of DAPI− CD45− CD31+ endothelial cells for mini-bulk RNA-seq (left). PCA showing global transcriptomic differences between endothelial cells from normal liver and malignant tissues (right). **F.** Heatmap showing decreased sinusoidal gene expression and increased capillarized gene expression in ECs isolated from malignant tissues compared with ECs isolated from adjacent normal liver. **G.** Immunofluorescence staining and quantification of lectin+ functional vessels in normal liver, breast cancer liver metastasis (LvMet), and orthotopic breast tumors (fat pad). Scale bar, 50 μm. **H.** Scanning electron microscopy (SEM) images showing the vascular wall structure (scale bar, 5 μm) and comparison of fenestration patterns between normal liver vessels and LvMet vessels. **I.** Immunostaining of laminin deposition and the sinusoidal marker Lyve1 (scale bar, 50 μm). Laminin deposition was quantified as the frequency of laminin+ VE-cadherin+ vessels among VE-cadherin+ cells in TECs of liver malignancies and NECs of adjacent normal (adj. N) liver. Lyve1 loss was quantified as the frequency of Lyve1+ VE-cadherin+ vessels among VE-cadherin+ vessels. **J.** Immunostaining of clinical breast cancer LvMet samples showing loss of the sinusoidal marker Lyve1 in TECs (scale bar, 50 μm). Representative patient samples are shown; breast cancer subtypes are labeled by color. Mean□±□s.e.m. shown. Smaller dots represent values from individual fields, and outlined circles indicate mean values taken over multiple fields/sections from the same mouse **(G-I)**. *P* values were calculated using two-tailed unpaired **(C, G, H, J)** and paired **(I)** Student’s *t*-test.

To validate the bioinformatic findings in human samples, we employed our previously established mouse models of liver malignancy: murine breast cancer cells (E0771) engrafted into the liver via portal vein injection, or hydrodynamic tail vein delivery of three plasmids to generate Myc-overexpressing, TP53-deleted hepatocellular carcinoma^28^. Once liver malignancies were established, we isolated fluorescence-activated cell sorting (FACS)-purified endothelial cells from liver malignancies and adjacent normal liver for bulk RNA-seq (**Fig. 1E**) and dynamic single-cell RNA-seq (Supplementary Fig. S1C), in which newly synthesized mRNA is labeled with the ribonucleoside analog 4-thiouridine (S4U) and identified by T-to-C conversions in sequenced reads^29^. Consistent with the human findings, TECs from both breast cancer liver metastases (LvMet) and HCC exhibited elevated expression of capillarized markers and reduced expression of sinusoidal markers (Fig. 1F, Supplementary Fig. S1D).

Beyond transcriptomic profiling, we sought to examine vascular abnormalities at the tissue level using the E0771 experimental LvMet model. For comparison, an orthotopic E0771 breast tumor was generated by fat pad injection. Vascular functionality was assessed using a fluorescence-labeled lectin perfusion assay^30^, in which circulating lectin binds endothelial glycoproteins only where blood flow is maintained, yielding fluorescent labeling of perfused vessels. Notably, LvMet-associated vessels exhibited significantly lower lectin binding than normal liver vessels, as expected, and even lower than primary tumor vessels, indicating profoundly dysfunctional vasculature in liver metastases (**Fig. 1G**).

A series of histological traits accompanied this vascular dysfunction in LvMet, including a significant reduction in fenestrae, with direct visualization revealing the loss of the small pores of LSECs (**Fig. 1H**), as well as pronounced laminin deposition and a progressive decrease in Lyve1 expression toward the tumor core (**Fig. 1I**). Consistent with the E0771 LvMet model, TECs in the autochthonous HCC model likewise exhibited enhanced laminin deposition, decreased Lyve1 expression, and reduced lectin perfusion (**Supplementary Fig. S1E,F**).

Importantly, these findings in mouse models were validated in patient samples from breast cancer LvMet (**Fig. 1J**) and HCC (**Supplementary Fig. S1G**), which similarly exhibited reduced vessel density and decreased expression of the sinusoidal marker LYVE1 in malignant tissues. Collectively, these results reveal profound disparities in the function, morphology, and transcriptional profiles of vessels between normal and malignant liver tissues. However, these phenotypic discrepancies raise two possibilities that require definitive lineage tracing to distinguish: (1) a vasculogenesis model, in which endothelial progenitor cells are recruited to the tumor and form Lyve1-negative vessels de novo; or (2) an angiogenic model, in which existing Lyve1+ LSECs infiltrate the tumor and undergo a sinusoidal-to-capillarized transition. Although the vessel co-option paradigm favors the latter, rigorous lineage tracing is required to discriminate between these models.

### Liver malignancy vessels originate from LSECs and grow by ingression

To elucidate the origin of TECs, we generated *Lyve1^Cre^;R26^LSL-tdT^*mice, in which an EGFP-Cre fusion protein is expressed under the control of the sinusoidal marker Lyve1. Cre-mediated recombination induces CAG promoter-driven irreversible expression of red fluorescent protein tdTomato at the Rosa26 locus (**Supplementary Fig. S2A**). Thus, even after Lyve1 silencing, as observed in TECs, cells derived from Lyve1+ LSECs remain tdTomato+, enabling their distinction from endothelial cells of non-LSEC origin, such as arterial endothelial cells or circulating endothelial progenitor cells. Although this *Lyve1^Cre^* line was originally generated to study the chemoattractant role of lymphatic vessels^31^, we found that the tdTomato+ cells were Lyve1+ but Prox1− in liver (**Supplementary Fig. S2B**), confirming that they are not lymphatic endothelial cells. Furthermore, flow cytometry showed that over 95% of endothelial cells in normal liver were tdTomato+, whereas fewer than 5% of CD45−CD31− cells were tdTomato+, demonstrating the efficiency and specificity of *Lyve1^Cre^* mice for labeling LSECs (**Supplementary Fig. S2B**). Notably, although approximately 15% of CD45+ cells were tdTomato+, likely reflecting Lyve1 expression in a subset of Kupffer cells^32,33^, this does not confound our analysis, as endothelial cells were gated as CD31+CD45−.

Remarkably, despite the significant reduction in Lyve1 expression in TECs, the percentage of tdTomato+ TECs remained unchanged in both LvMet and HCC, as demonstrated by immunostaining (**Fig. 2A**, **Supplementary Fig. S2D**) and flow cytometry quantification (**Fig. 2B**, **Supplementary Fig. S2E**). These findings strongly support the angiogenesis model that TECs in liver malignancies originate from pre-existing LSECs of the normal liver.

**Figure 2.**
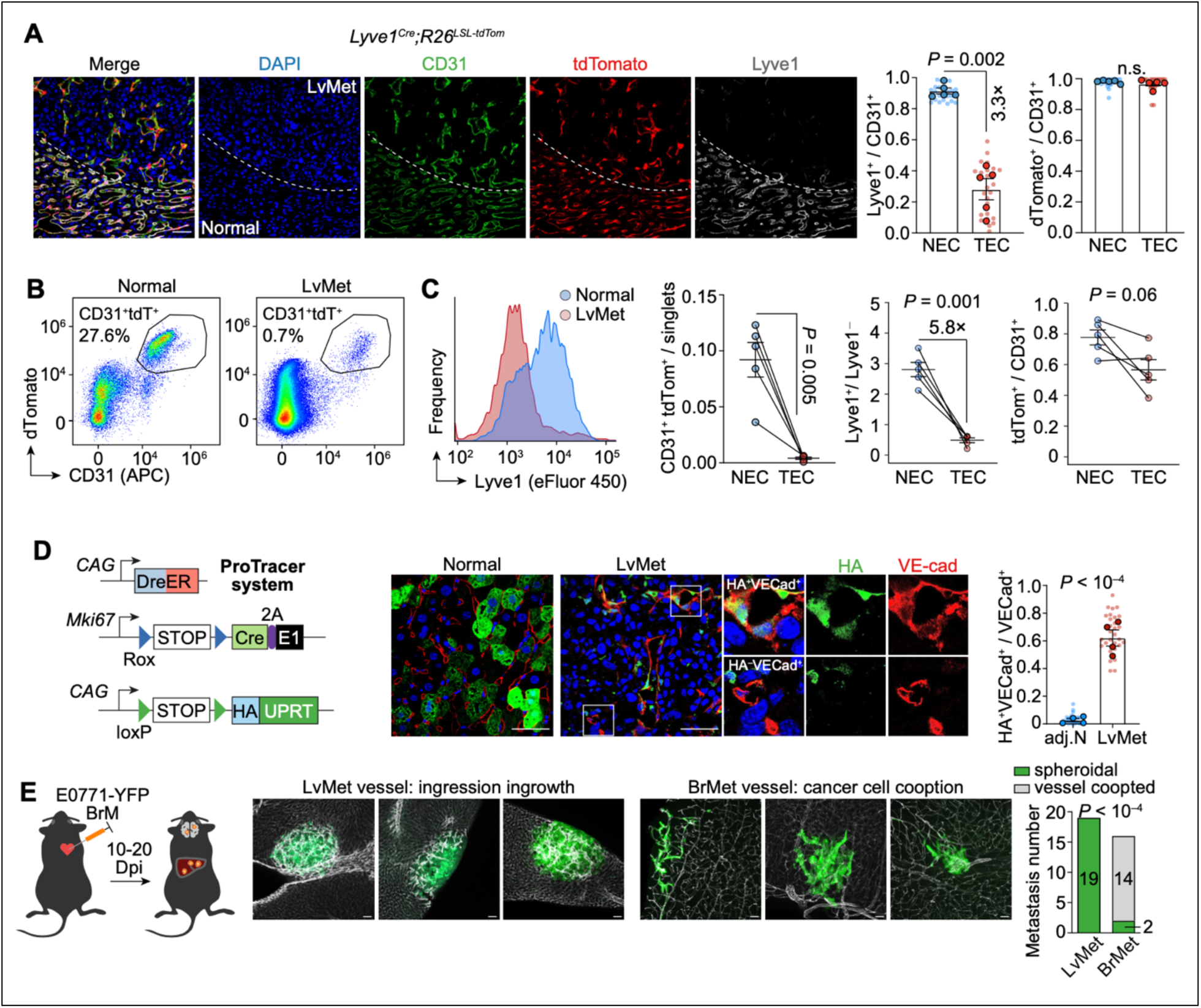
Lineage tracing of LSEC in breast cancer liver metastasis. **A.** Immunostaining (left) and quantification of Lyve1 and tdTomato in CD31+ NECs (endothelial cells in adjacent normal tissue) and CD31+ TECs (endothelial cells in breast cancer liver metastasis) in the Lyve1 lineage-tracing system (*Lyve1^Cre^; R26^LSL-tdTomato^*) (right). Scale bar, 50 μm. **B.** Flow cytometry plot showing the gating strategy for tdTomato+ CD31+ endothelial cells. **C.** Histogram showing expression of the sinusoidal marker Lyve1 in CD45− CD31+ TECs and CD45− CD31+ NECs (left). Flow cytometric quantification showing decreased Lyve1 expression and reduced endothelial cell abundance, while the percentage of tdTomato+ cells among CD31+ endothelial cells remained unchanged (right). **D.** Proliferation tracing system (H11^DreER^; Ki67^RSR-Cre^; R26^LSL-HA-UPRT^) revealing an increased percentage of HA-tag+ proliferating TECs in LvMet, despite lower vascular density. **E.** Comparison of incipient metastasis shapes between liver metastasis (LvMet) and brain metastasis (BrMet), suggesting distinct endothelial growth patterns. Mean□±□s.e.m. shown. Smaller dots represent values from individual fields, and outlined circles indicate mean values taken over multiple fields/sections from the same mouse **(A, D)**. *P* values were calculated using two-tailed paired Student’s *t*-test (**A, C, D**) and Fisher’s exact test (**E**).

In addition to Lyve1-based lineage tracing, we employed a proliferation tracing system^34^ that we previously used to reveal local proliferation of LvMet macrophages^28^ and to label proliferating PyMT cells^35^. In this model, cells that undergo proliferation within a defined time window are permanently labeled with a human influenza hemagglutinin (HA)-tagged protein. Strikingly, immunostaining for the HA tag showed that vessels in adjacent normal liver were almost entirely HA-negative, whereas LvMet-associated vessels were mostly HA-positive, indicating that LvMet vessels had undergone at least one round of proliferation.

As vessel co-option is also exploited by incipient brain metastases (BrMet) of triple-negative breast cancer and lung adenocarcinoma^36^, we sought to examine whether the growth patterns are similar to or distinct from those in liver metastasis. Because parental E0771 cells poorly colonize the brain after intracardiac injection, we first performed two rounds of in vivo selection^37^ to enrich for E0771 variants with high brain-colonizing propensity (denoted E0771-BrM). We harvested brains of mice injected intracardially with E0771-BrM and livers of mice injected intraportally with parental E0771, both labeled with yellow fluorescent protein (YFP), at 10-20 days after injection to trace incipient metastatic lesions smaller than 1 mm in diameter. Interestingly, the majority of E0771 BrMet lesions exhibited a vascular wrapping pattern, consistent with a previous report^36^. In contrast, even very small liver metastatic lesions exhibited a spheroidal shape. We therefore propose that E0771 adopts distinct growth patterns in different organs, even though cancer cells exploit pre-existing vasculature in both settings: in the brain, extravasated cancer cells slide and grow along the vessels, whereas in the liver, LSECs protrude inward into the LvMet lesion, a growth pattern we term “ingression”.

### Lyve1 restoration fails to normalize TECs and exacerbates capillarization

Next, we investigated whether the loss of the sinusoidal marker Lyve1 contributes to the abnormalities observed in TECs. Inspired by a study showing that restoring LSEC differentiation can reverse cirrhosis^23^, we hypothesized that re-expression of Lyve1 in TECs may restore the sinusoidal features of LvMet vessels. To test this hypothesis, we developed two Cre-inducible Lyve1 overexpression models. The first model employs a doxycycline (Dox)-inducible, Cre-dependent allele targeted to the TIGRE safe-harbor locus^38^ (**Supplementary Fig. S3A**), denoted as Lyve1^OE1^. In this model, Cre recombinase induces expression of the reverse tetracycline-controlled transactivator (rtTA); in the presence of Dox, rtTA drives expression of Flag-tagged Lyve1, allowing it to be distinguished from endogenous, non-tagged Lyve1. After establishing the E0771 experimental liver metastasis model in Lyve1^OE1^ mice and littermate controls lacking the TIGRE knock-in allele, we administered Dox to induce ectopic Lyve1 expression in TECs. Flag-tagged Lyve1 was successfully expressed in both normal liver and LvMet tissues of Lyve1^OE1^ mice (**Supplementary Fig. S3B,C**). However, significantly more Flag+ than tdTomato+ cells were observed (**Supplementary Fig. S3D**), which we attribute to leakiness of the tetracycline response element (TRE) promoter, driving expression even in the absence of rtTA.

To circumvent nonspecific Lyve1 expression in non-LSEC-derived cells, we generated a second model, denoted as Lyve1^OE2^, in which ectopic expression of Flag-tagged Lyve1 is controlled solely by Cre, resulting in Lyve1 upregulation specifically in cells that already express Lyve1 (**Fig. 3A**). We then injected E0771 cells engineered to express nuclear-localized H2B-HaloTag into the portal vein to establish LvMet, enabling delineation of the LvMet-normal liver border. Consistent with the Dox-inducible Lyve1^OE1^ model, Lyve1^OE2^ successfully restored Lyve1 expression in TECs in LvMet (**Fig. 3B,C**). Importantly, Flag expression was restricted to VE-cadherin+ TECs, resolving the issue of nonspecific expression.

**Figure 3.**
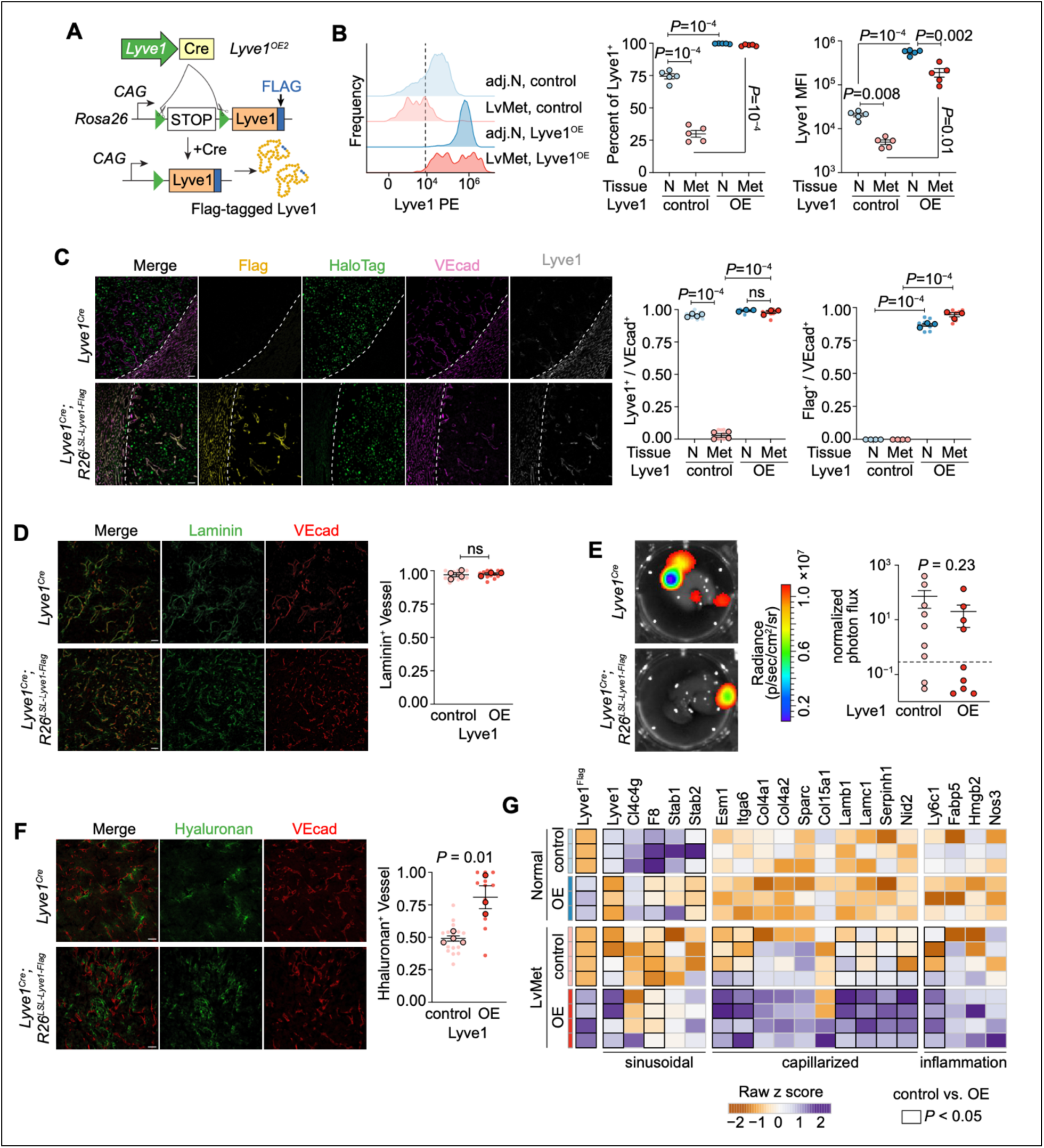
TEC-specific Lyve1 restoration failed to reverse sinusoidal-to-capillarized transition. **A.** Schematic of the endothelial-specific conditional Lyve1 overexpression (Lyve1^OE2^) model (genotype: *Lyve1^Cre^; R26 ^Lyve1-Flag^*). **B,C.** Flow cytometry **(B)** and immunostaining **(C)** of Lyve1 in Lyve1^OE2^ mice and littermate controls (scale bar, 50 μm). Ectopic Flag-tagged Lyve1 expression not only restored Lyve1 expression in Lyve1-low TECs but also increased Lyve1 expression in already Lyve1+ NECs. **D.** Restoration of Lyve1 in TECs failed to decrease vascular laminin deposition, as shown by immunofluorescence staining (scale bar, 50 μm). **E.** Metastatic outgrowth was similar between Lyve1^OE2^ and littermate control mice. Bioluminescence-negative samples fall below the threshold (dashed line). **F.** Ectopic Lyve1 expression unexpectedly increased hyaluronan deposition in the tumor microenvironment. **G.** Heatmap of RNA-seq data (Log2(TPM+1)) from TECs and NECs isolated from LvMet and adjacent normal liver. Ectopic Lyve1 expression decreased endogenous Lyve1 expression as well as other sinusoidal markers. Simultaneously, capillarized gene expression in TECs increased in Lyve1^OE2^ mice. Mean□±□s.e.m. shown. Smaller dots represent values from individual fields, and outlined circles indicate mean values taken over multiple fields/sections from the same mouse **(C, D, F)**. *P* value was calculated using two-tailed unpaired Student’s *t*-test **(B-D, F)**.

We then evaluated the effect of Lyve1 restoration on TEC phenotypes and metastatic outgrowth. Given the high variability of tumor growth in the autochthonous HCC model, we used the E0771 experimental liver metastasis model for all subsequent functional perturbations. Laminin deposition remained unchanged (**Fig. 3D**), suggesting persistence of the capillarized phenotype. Furthermore, metastatic outgrowth did not differ significantly between Lyve1^OE2^ mice and controls (**Fig. 3E**). Paradoxically, hyaluronan, the extracellular matrix (ECM) glycosaminoglycan endocytosed by Lyve1, was augmented in LvMet of Lyve1^OE2^ mice (**Fig. 3F**). To resolve this paradox, we performed RNA-seq on FACS-purified LSECs isolated from LvMet and adjacent normal liver of Lyve1^OE2^ mice and littermate controls. Ectopic Lyve1 expression imposed a compensatory response: endogenous Lyve1 expression was decreased, along with other scavenger receptors such as F8 and Stab2, in normal liver LSECs (**Fig. 3G**). Notably, Stab2, together with Lyve1, mediates endocytic clearance of hyaluronan^11,20,39^, such that the concerted downregulation of Stab2 may explain the paradoxical hyaluronan accumulation in LvMet of Lyve1^OE2^ mice.

Several ECM proteins marking capillarized features in TECs were also upregulated (**Fig. 3G**). Collectively, these results indicate that Lyve1 restoration cannot revert the sinusoidal-to-capillarized transition, which is governed by additional regulators beyond Lyve1. On the contrary, the capillarized phenotype was exacerbated, possibly because ectopic Lyve1 suppresses endogenous scavenger receptor expression, thereby attenuating sinusoidal features.

### Inflammation restrains angiogenesis in liver metastasis of breast cancer

In addition to the loss of sinusoidal features, heightened inflammation is a second characteristic of TECs in liver malignancies^20,40^. To explore the functional consequences of inhibiting key inflammatory signaling pathways, we first profiled publicly available spatial transcriptomic data from four human HCC and two human colorectal cancer liver metastasis samples to nominate candidate pathways. Hematoxylin and eosin (H&E) staining was used to delineate normal liver, tumor, and border regions (**Supplementary Fig. S4A**). We selected 11 cytokines that have been implicated in a broad range of chronic liver diseases^41–47^. Notably, most of these cytokines were elevated at the normal liver–carcinoma border (**Supplementary Fig. S4A**), with CCL2 and TNF being the most striking, showing this pattern in all patients.

To validate these observations in human samples, we employed Ccr2-GFP reporter mice and Tnfr1 knockout mice to probe the roles of CCL2-CCR2 and tumor necrosis factor α (TNFα) signaling in tumor angiogenesis, respectively. In this reporter allele, the coding region of Ccr2 is replaced by green fluorescent protein (GFP); thus, heterozygous mice serve as fluorescent reporters of CCR2-expressing inflammatory mononuclear phagocytes, including monocytes and macrophages, whereas homozygous mice lack functional CCR2, functionally ablating the CCL2-CCR2 axis. Consistent with the clinical observations, GFP+ inflammatory mononuclear cells were markedly enriched at the tumor border, with numbers declining toward the tumor core (**Fig. 4A**). Ablation of Ccr2 significantly decreased the number of GFP+ cells in both the border and the tumor core, which, interestingly, resulted in a significant increase in vascular density at the border, suggesting that CCL2-driven inflammation normally restrains vascular density at the border. Two points merit emphasis. First, even in the Ccr2-deficient background, GFP+ cells remained enriched at the border, indicating that CCL2-CCR2-independent inflammatory signals also recruit these cells. Second, Ccr2 ablation had no effect on angiogenesis in the tumor core, suggesting that compensatory mechanisms counterbalance the restraining effect of inflammation on vascular density in the core.

**Figure 4.**
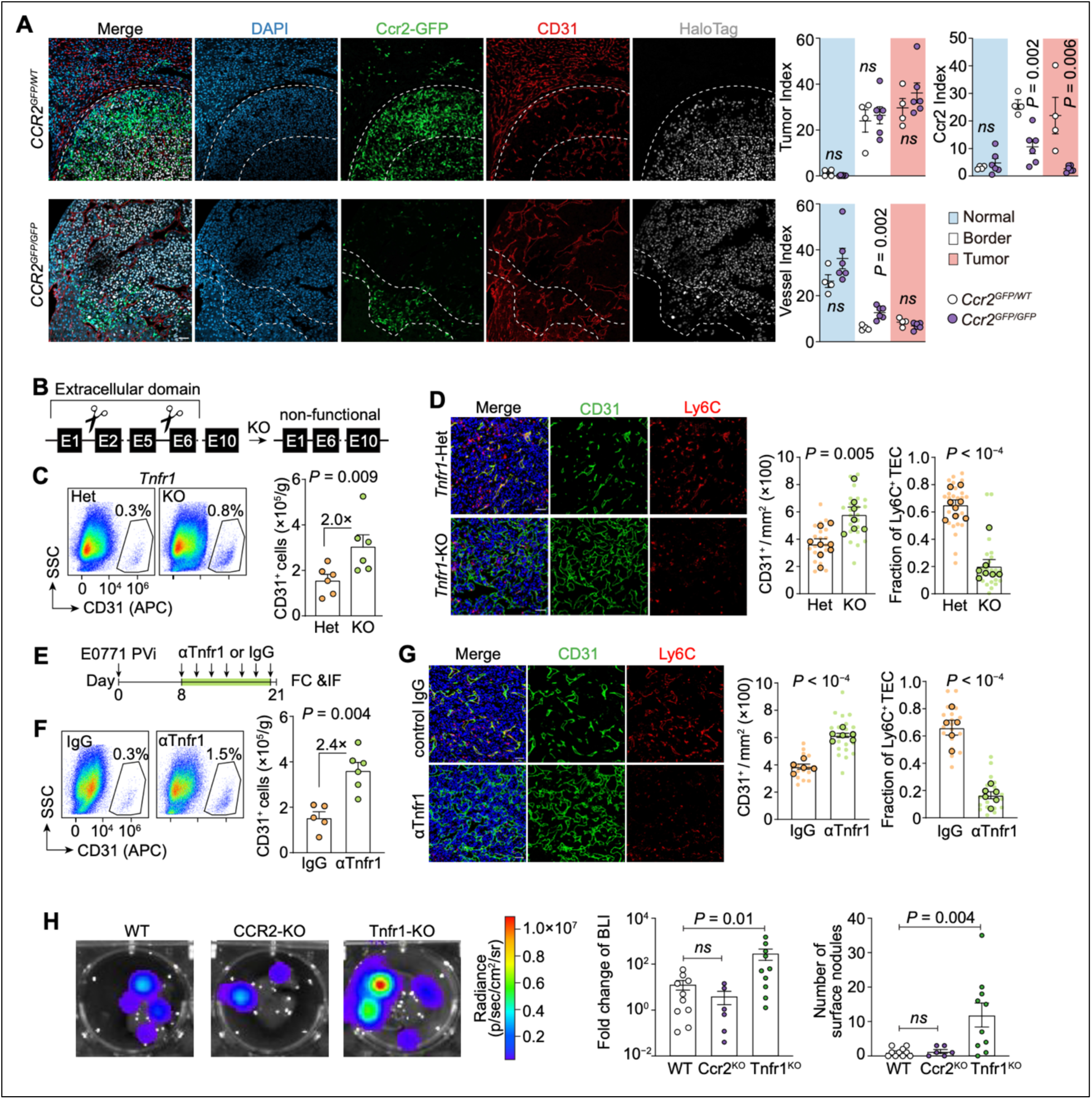
Blocking CCR2 or TNF signaling reduces vascular inflammation while promoting angiogenesis in TECs. **A.** Immunofluorescence quantification of CCR2+ inflammatory monocyte infiltration and vascular density in adjacent normal liver, the LvMet area (as defined by E0771 cells engineered to express nuclear-localized H2B-HaloTag), and the border area. Tumor, CCR2, and vessel indices were calculated by dividing the HaloTag+, GFP+, and CD31+ areas (pixels), respectively, by the number of DAPI+ nuclei in each area. Scale bar, 50 μm. **B.** Schematic of Tnfr1 knockout (KO) mice, showing deletion of the extracellular domain responsible for sensing TNFα. **C.** Flow cytometry quantification of CD31+ TEC number in LvMet from Tnfr1 KO mice and heterozygous littermate controls (Het). **D.** Immunostaining quantification of vessel density (left) and vascular expression of the inflammatory marker Ly6C, quantified as the fraction of Ly6C+ TECs (right), in TECs from LvMet of Tnfr1 KO mice and heterozygous littermate controls (Het). **E.** Schematic of the experimental design for antibody-mediated blockade of Tnfr1. Mice were intraperitoneally injected with 100 μg anti-Tnfr1 antibody or an equivalent amount of isotype control antibody every other day starting from Day 8 after portal vein injection, with treatment lasting 2 weeks (IF, immunofluorescence; FC, flow cytometry). **F.** Flow cytometry quantification of CD31+ TEC number in LvMet from mice treated with anti-Tnfr1 antibody or isotype control antibody. **G.** Immunostaining quantification of vessel density (left) and vascular Ly6C expression (right) in TECs from LvMet of mice treated with anti-Tnfr1 antibody or isotype control antibody (scale bar, 50 μm). **H.** Bioluminescence imaging (BLI) quantifying the metastatic outgrowth rate in wild-type, Ccr2 KO, and Tnfr1 KO backgrounds. Fold change in BLI was calculated by dividing the final ex vivo liver BLI intensity by the Day 0 in vivo BLI intensity. Mean□±□s.e.m. shown. Smaller dots represent values from individual fields, and outlined circles indicate mean values taken over multiple fields/sections from the same mouse **(D, G)**. *P* values were calculated using two-tailed paired Student’s *t*-test **(A)** and unpaired Student’s t-test (C, D, F–H).

In addition to CCL2-CCR2 signaling, we next examined TNF signaling, which has been implicated in both angiogenesis and inflammation^48^. TNFα signals through two major receptors: Tnfr1, which is ubiquitously expressed and binds both soluble and membrane-bound TNFα, and Tnfr2, which is primarily restricted to immune cells and binds only membrane-bound TNFα^49^. The broader binding capacity of Tnfr1 makes it a more versatile target for modulating TNF signaling^50^. We first employed Tnfr1 knockout (KO) mice to examine the role of Tnfr1 in mediating inflammatory responses in TECs within LvMet. In this knockout line, the extracellular domain of Tnfr1 is absent, rendering the receptor non-functional and incapable of sensing TNFα (**Fig. 4B**). Tnfr1 knockout significantly increased vessel density, as shown by both flow cytometry (**Fig. 4C**) and immunofluorescence staining (**Fig. 4D**). Remarkably, vascular Ly6C expression, a classical inflammatory monocyte marker whose elevation has also been reported in various vascular inflammatory conditions including atherosclerosis^51^, pancreatic cancer^52^ and colorectal cancer^53^, was profoundly decreased (**Fig. 4D**). To corroborate the findings in Tnfr1 KO mice, we employed a blocking anti-Tnfr1 antibody (**Fig. 4E**), which enables time-controlled blockade of Tnfr1 signaling (e.g., at the beginning of micrometastasis formation). Consistent with the phenotypes observed in Tnfr1 KO mice, antibody-mediated blockade of Tnfr1 promoted angiogenesis and reduced Ly6C expression in TECs (**Fig. 4F,G**).

Because inhibiting inflammatory signaling may dampen the inflammatory status of TECs while enhancing angiogenesis, two phenotypes with potentially opposite effects on metastasis, we sought to determine whether blockade of Ccr2 or Tnfr1 modulates metastatic outgrowth. We quantified metastatic burden by bioluminescence imaging and by counting nodules on the liver surface. Although metastatic burden remained similar between wild-type controls and Ccr2 knockout mice, consistent with previous reports^54^, Tnfr1 knockout significantly potentiated metastatic outgrowth (**Fig. 4H**). Collectively, these results highlight a dual role of inflammation in tumor microenvironment: it may potentially drive vascular dysfunction on one hand, yet restrains angiogenesis on the other. Consequently, inhibition of inflammatory signaling, such as Tnfr1 blockade, enhanced metastatic burden, possibly because the resulting angiogenesis provides additional vascular support for the growth and spread of metastatic cancer cells within the liver.

### Epigenetic reprogramming underlies the capillarized state of TECs

To understand the mechanisms underlying the sinusoidal-to-capillarized transition of TECs, we crossed the Lyve1^Cre^ line with NuTRAP mice for in situ multi-omic profiling (**Fig. 5A**). In this system, Cre-expressing cells express an EGFP-conjugated ribosomal subunit (L10a) that labels both the nucleus and translating mRNA^55^. Because TECs are rare in the tumor microenvironment and prolonged FACS purification can perturb the epigenetic landscape, NuTRAP enables rapid tissue lysis and nuclear fixation before fluorescence-activated nuclear sorting, preserving the epigenetic state (**Fig. 5B,C**). Immunofluorescence confirmed efficient GFP labeling of vessels in both LvMet regions (HaloTag+) and adjacent normal tissue, validating this model for *in situ* epigenetic profiling of TECs (**Fig. 5D**).

**Figure 5.**
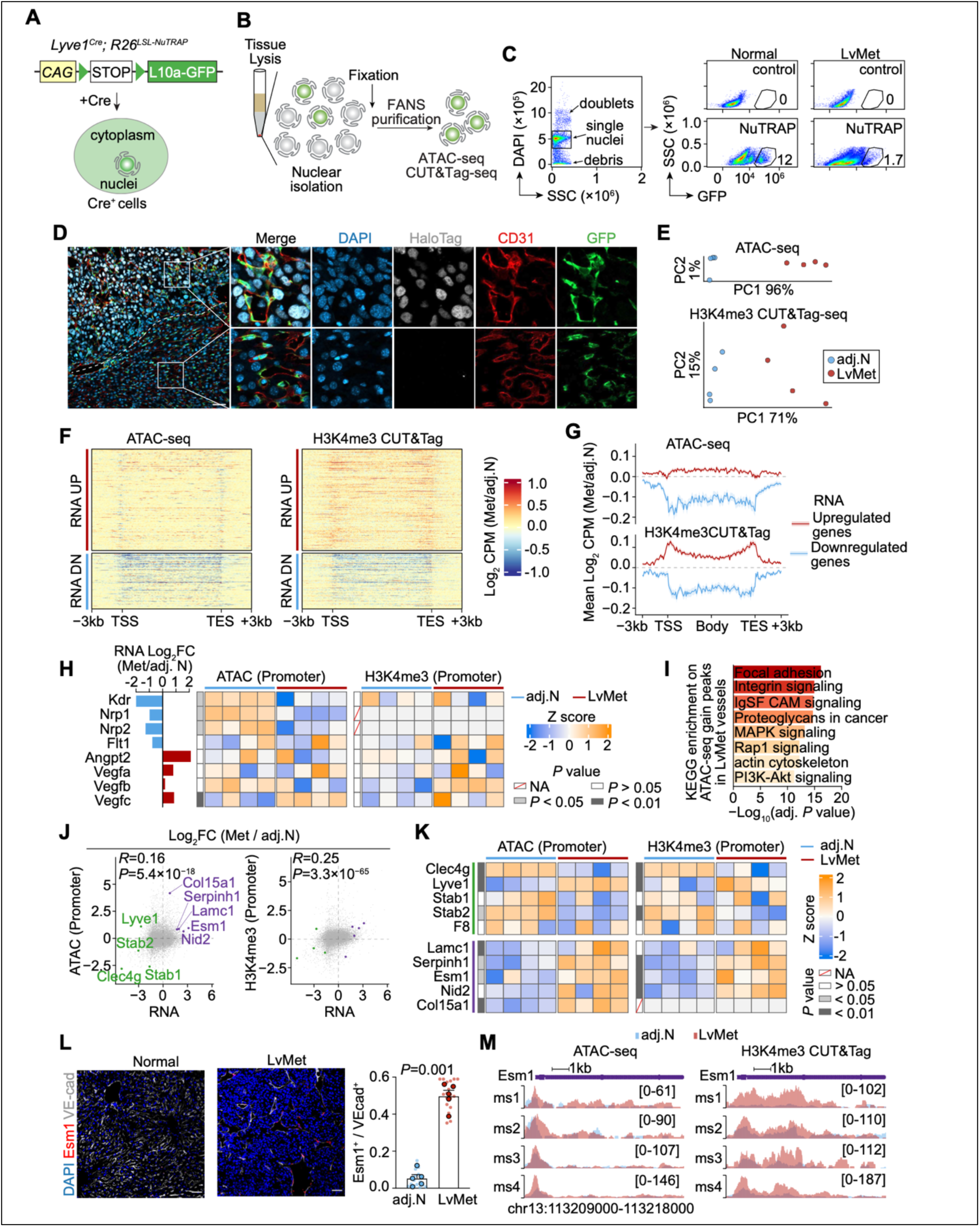
Epigenetic reprogramming underlies the sinusoidal-to-capillarized transition. **A,B.** Schematic of the endothelial-specific NuTRAP mouse model *Lyve1^Cre^; R26 ^LSL-L10a–EGFP^* (**A**) and experimental workflow for nuclei isolation for epigenetic profiling (**B**). **C.** Because ribosome assembly occurs in the nucleolus, the EGFP-tagged ribosomal protein L10a labels intact nuclei, allowing flow cytometric sorting of GFP+ nuclei from normal liver and LvMet tumors after fixation. **D.** Immunostaining showing the endothelial specificity of NuTRAP labeling in both LvMet and adjacent normal (adj. N) liver. **E.** Principal-component analysis of the epigenetic landscape of endothelial cells isolated from LvMet and adj. N liver, computed on variance-stabilized counts over all consensus peaks. The contributions of PC1 and PC2 are indicated on the axes. **F.** Metagene heatmaps of changes in chromatin accessibility (ATAC-seq) and H3K4me3 (CUT&Tag) between LvMet-associated and adj. N liver vessels across the entire gene locus for the top 500 RNA-upregulated and RNA-downregulated genes (nominal *P* < 0.05, |log2FC| > 1, ranked by *P*). Regions span 3 kb upstream of the transcription start site (TSS), the scaled gene body, and 3 kb downstream of the transcription end site (TES). **G.** Profile plot corresponding to (**F**), showing mean signals with standard error bands for RNA-upregulated and RNA-downregulated genes. **H.** Promoter signal (Z-scored Log2 CPM) heatmap of the eight VEGF-axis genes for ATAC-seq (left) and H3K4me3 (right). RNA Log2FC of each gene is shown as the bar plot at left. **I.** KEGG pathway analysis of genes assigned to accessibility-gained peaks in tumor-associated endothelial cells (TECs) versus normal endothelial cells (NECs). **J.** Scatter plots of RNA Log2FC (TEC vs NEC) against promoter Log2FC (TEC vs NEC) for chromatin accessibility (left) and H3K4me3 (right). Each grey dot is one gene; selected sinusoidal (green) and capillarized (purple) genes are highlighted. **K.** Promoter signal (Z-scored Log2 CPM) heatmap of selected sinusoidal and capillarized marker genes for ATAC-seq (left) and H3K4me3 (right). **L.** Immunostaining and quantification of perivascular ESM1 expression in LvMet and adj. N liver (scale bar, 50 μm). **M.** Genome browser tracks of ATAC-seq and H3K4me3 CUT&Tag signals at the *Esm1* locus in paired normal and tumor vessels, showing gained signal at the promoter.

To assess the quality of the epigenetic profiling, we first examined concordance between the two modalities. Because both chromatin accessibility and the activating histone mark H3K4me3 are associated with gene activation, their promoter-level signals correlated well (**Supplementary Fig. S5A**). Moreover, genes epigenetically activated or repressed in TECs (**Supplementary Fig. S5B**) showed correspondingly increased or decreased RNA expression (**Supplementary Fig. S5C**), confirming the reliability of the profiling. Having established this, we found that TECs in LvMet and LSECs in adjacent normal liver exhibited marked differences in both chromatin accessibility and H3K4me3 modification, as determined by ATAC-seq and CUT&Tag-seq, respectively (**Fig. 5E-G**). Pathway analysis further revealed differential oxygen-response signaling (**Supplementary Fig. S5D**), consistent with the higher hypoxia levels in the tumor microenvironment.

Because VEGF signaling is widely regarded as the dominant driver of angiogenesis, we first examined VEGF receptors (Kdr, Nrp1, Nrp2, Flt1), ligands (Vegfa, Vegfb, Vegfc), and the sprouting angiogenesis marker Angpt2 at both transcriptomic and epigenetic levels. Chromatin accessibility of these genes was decreased in TECs, concordant with their reduced RNA expression (**Fig. 5H**). Thus, despite elevated Vegfa and Vegfc, reduced endothelial responsiveness suggests a limited contribution of VEGF signaling to the transition.

KEGG enrichment analysis of regions with gained chromatin accessibility revealed focal adhesion, integrin, and proteoglycan pathways (**Fig. 5I**), all linked to ECM deposition during vascular capillarization. We then examined the sinusoidal and capillarized marker genes defined earlier: sinusoidal genes (Clec4g and Stab2) were decreased in TECs at all levels of RNA, chromatin accessibility, and H3K4me3, whereas ECM-associated capillarized genes (Lamc1, Serpinh1, Esm1) were increased at all three levels (**Fig. 5J,K**). Notably, although Lyve1 RNA was decreased, its chromatin accessibility and H3K4me3 were increased, suggesting that Lyve1 downregulation is imposed post-transcriptionally rather than through epigenetic silencing.

We focused on Esm1 (endothelial cell-specific molecule 1) for two reasons: unlike other ECM proteins, it is an endothelial glycoprotein, and, as its name suggests and immunofluorescence confirmed (**Fig. 5L**), its expression is restricted to a subset of endothelial cells. Paired comparison of TECs and adjacent normal LSECs from the same mice showed that, although both chromatin accessibility and H3K4me3 increased at the Esm1 locus in TECs, the H3K4me3 gain was more pronounced; the already-open chromatin in normal LSECs suggests that Esm1 is poised for rapid activation in TECs (**Fig. 5L,M**).

### Targeting Esm1 promotes vascular normalization in breast cancer liver metastasis

Pseudotime analysis of our integrated scRNA-seq dataset showed that Esm1 gradually increased as sinusoidal vessels transdifferentiated into capillarized vessels, peaked in capillarized vessels, and decreased sharply in arterial endothelial cells (Supplementary Fig. S6A–D). In addition to serving as a capillarized marker, Esm1 has been reported to be critical for endothelial tip cell function during vascular migration^56^. To contextualize ESM1 relative to other sprouting angiogenesis markers, we also analyzed a publicly available pan-cancer scRNA-seq dataset comprising 116,958 vascular endothelial cells from 437 tumor samples^27^. ESM1 expression was the most restricted to tip cells, compared with ANGPT2 and VEGFA (Supplementary Fig. S6E). Pseudotime analysis further revealed distinct expression dynamics, with ESM1 peaking at the Tip S3 stage (Supplementary Fig. S6F,G), a phase during which tip cells undergo phenotypic conversion to integrate into arterioles. This link between ESM1 expression and phenotypic conversion pinpoints a role for ESM1 in endothelial cell plasticity and parallels the sinusoidal-to-capillarized transition of liver tumor vessels. Consistently, pan-cancer survival analysis in TCGA^57^ (Supplementary Fig. S6H), showed that high ESM1 expression is associated with poor prognosis, positioning ESM1 as a promising target for vascular normalization (VN) therapy.

To functionally probe the role of Esm1 in TECs, we generated *Lyve1^Cre^;Esm1^fl/fl^* (Esm1 cKO) mice (**Fig. 6A**). Although Lyve1 is expressed in both LSECs and lymphatic vessels, Esm1 expression in these cell types is negligible (**Fig. 5L**, **Fig. 6B**), ensuring that the observed effects of Esm1 cKO are specific to TECs and minimizing confounding effects on LSECs or lymphatic endothelium. Esm1 cKO led to a significant reduction in perivascular ESM1 levels and a modest decrease in the total number of TECs, as quantified by flow cytometry and immunostaining (**Fig. 6C,D**). Interestingly, in line with our earlier finding that Tnfr1 blockade increases vessel density (**Fig. 4D,G**), perivascular ESM1 deposition was elevated in LvMet vessels of Tnfr1 KO mice (**Fig. 6E**). This suggests that the increased vessel density in LvMet of Tnfr1 knockout mice may be partially attributed to ESM1, which counteracts the pro-apoptotic effects of TNF-driven inflammation.

**Figure 6.**
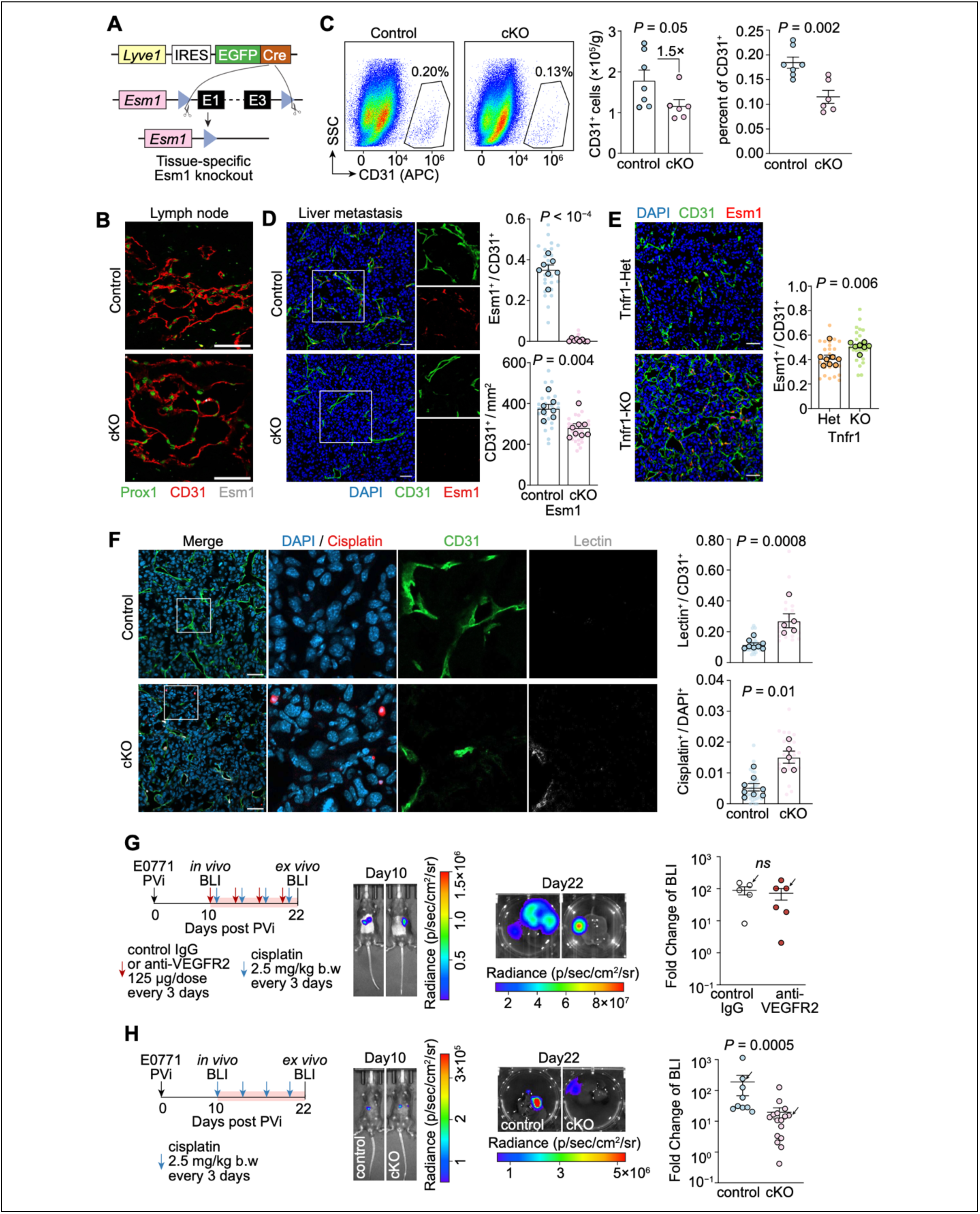
Targeting Esm1 normalizes vessels in breast cancer liver metastasis. **A.** Schematic of conditional knockout (cKO) of Esm1 upon exposure to Cre recombinase. **B.** Immunostaining of ESM1 in lymphatic vessels from Esm1 cKO mice and littermate controls. Esm1 deletion does not alter lymphatic vessel morphology, as Esm1 is not expressed in lymphatic vessels (scale bar, 50 μm). **C.** Flow cytometry quantification of the absolute number of CD31+ TECs and the percentage of CD31+ TECs in Esm1 cKO mice and littermate controls. **D.** Immunofluorescence quantification of perivascular ESM1 expression and the effect of Esm1 knockout on vessel density (CD31+ vessels/mm²) in LvMet. **E.** Immunostaining showing the effect of Tnfr1 knockout (KO) on vessel density and perivascular ESM1 deposition (scale bar, 50 μm). **F.** Immunostaining quantification of vascular lectin perfusion efficiency (lectin+/CD31+) and short-term cisplatin delivery (cisplatin+/DAPI+) in LvMet of Esm1 cKO and littermate controls. For short-term treatment, cisplatin was intraperitoneally injected at 10 mg/kg body weight 6 hours before tumor dissection. **G,H.** Bioluminescence imaging (BLI) quantification of metastatic burden to assess the effect of VEGFR inhibition (**G**) and endothelial-specific Esm1 ablation (**H**) on cisplatin efficacy in inhibiting metastatic outgrowth. Mice were bioluminescence-imaged at day 10 after portal vein injection, of which the BLI intensity was used to normalize the ex vivo liver metastatic signal at the endpoint to calculate the BLI fold change. Mice used for representative images are indicated by arrowheads. Mean□±□s.e.m. shown. Smaller dots represent values from individual fields, and outlined circles indicate mean values taken over multiple fields/sections from the same mouse **(D, E, F)**. *P* values were calculated using two-tailed unpaired Student’s *t*-test **(C-H)**.

We then examined the effect of Esm1 cKO on VN and its potential to synergize with chemotherapy in inhibiting metastatic outgrowth. We employed a short-term, high-dose cisplatin regimen (10 mg/kg) combined with lectin perfusion in Esm1 cKO mice and littermate controls bearing LvMet. Immunostaining for cisplatin-DNA adducts and fluorescently labeled lectin showed that Esm1 cKO improved both vessel perfusion and chemotherapeutic delivery despite reduced vessel density (**Fig. 6F**), two defining features of vascular normalization.

Because twice-weekly high-dose cisplatin caused body weight loss, we instead used a lower dose (2.5 mg/kg) every three days for long-term treatment to examine the effect of Esm1 cKO-induced VN on metastatic outgrowth. As a comparison, we combined cisplatin with a blocking anti-VEGFR2 antibody. After micrometastases were established, as inferred from bioluminescence imaging, mice were randomized to receive anti-VEGFR2 antibody or IgG isotype control. Consistent with the limited VEGF contribution suggested in **Fig. 5H**, where Kdr, the gene encoding VEGFR2, was decreased at both RNA and chromatin accessibility levels, VEGFR2 inhibition did not alter metastatic outgrowth (**Fig. 6G**). In contrast, Esm1 cKO significantly decreased metastatic burden by 10-fold (**Fig. 6H**), suggesting that Esm1 cKO-induced VN enhances chemotherapeutic efficacy. Collectively, these data establish Esm1 as a target for promoting VN and synergizing with chemotherapy in liver metastasis.

## Discussion

The present study identifies key characteristics of vessels in liver malignancies, including breast cancer LvMet and HCC. Similar to the pathogenesis of cirrhosis, vessels in liver malignancies lose their sinusoidal features and undergo capillarization^23^. To elucidate the phenotypic plasticity underlying this process, we employed a *Lyve1^Cre^*lineage-tracing system. Commonly used endothelial-specific Cre lines, such as *VE-cadherin^Cre^*, are unsuitable for this purpose, as VE-cadherin is expressed in both LSECs and TECs^58^, and therefore cannot distinguish their lineages. In contrast, the *Lyve1^Cre^* model reveals that Lyve1+ LSECs can give rise to Lyve1− TECs, providing definitive evidence for the origin of TECs in liver malignancies. Notably, although cancer cells in the liver have been proposed to co-opt normal liver vasculature^8,14^, our findings reveal that this co-option differs fundamentally from that in brain metastasis: liver metastases grow spheroidally and attract the ingression of LSECs, whereas brain metastatic cells grow along and wrap around pre-existing vessels.

An intriguing observation from our proliferation lineage-tracing experiments is that, although TECs in LvMet have undergone at least one round of proliferation, vascular density within LvMet remains lower than that of adjacent normal liver. Our Tnfr1 knockout model may reconcile this paradox. Pro-angiogenic cues in the tumor microenvironment may drive TEC proliferation, while TNF signaling, which exerts both pro-inflammatory and pro-apoptotic effects, may simultaneously impose vascular trimming. The net result is a vasculature that is proliferative yet sparse, proliferating to support tumor ingrowth, but continuously pruned by inflammatory apoptosis, and therefore unable to reach homeostatic density.

Another important concept emerging from our study is that the correlated phenotypes observed in TECs of LvMet do not necessarily causally reinforce one another, and targeting a single feature can even exacerbate the others. For example, restoring Lyve1 unexpectedly augmented extracellular matrix (ECM) deposition, because ectopic Lyve1 expression suppressed endogenous Lyve1 and other sinusoidal genes, thereby exacerbating the capillarized phenotype. Similarly, blockade of TNF signaling enhanced Esm1 expression, thereby promoting angiogenesis and capillarization. These compensatory responses indicate that the vascular abnormalities in liver metastasis form an interconnected network rather than a list of independent targets, and that effective normalization strategies must account for such coupling.

Because vascular normalization based on targeting VEGF signaling has limited efficacy in treating liver malignancies^59^, we propose targeting the ECM protein ESM1 as an effective approach for normalizing TECs in LvMet. Our in situ epigenetic profiling provides the mechanistic basis: chromatin accessibility of VEGF receptors is decreased in TECs, whereas the Esm1 locus shows increased accessibility and H3K4me3. Notably, some accessibility is already present in normal LSECs, rendering Esm1 poised for rapid activation in the LvMet microenvironment.

Several properties position ESM1 as a compelling target. First, it unites the two programs of the capillarized state: it is an ECM protein deposited by capillarized vessels and a tip cell marker that triggers sprouting angiogenesis. Second, its expression is TEC-restricted: ESM1 signal was confined to the vicinity of TECs, limiting off-tumor effects. Third, although ESM1 has been shown to promote angiogenesis by increasing VEGFA bioavailability^60^, our data show that ESM1 promotes angiogenesis even in VEGF-low-responsive TECs, indicating a VEGF-independent function.

Translationally, our Lyve1-lineage cKO spared LSECs and lymphatic vessels, which express little Esm1, and accordingly showed no observed toxicity. However, pharmacological Esm1 blockade would act systemically, and its toxicity requires careful evaluation. Moreover, the targeting strategy itself warrants consideration: ESM1 is an O-linked glycoprotein whose Ser156 glycosylation is important for its growth factor mitogenic activity^61^.

In summary, our study reveals the origin, phenotypic plasticity, and pathological features of TECs in liver malignancies, and highlights ESM1 blockade as a promising strategy for normalizing dysfunctional vessels. These findings offer new therapeutic avenues, including potential synergy with chemotherapy, for improving vascular function in liver malignancies and, given the pan-cancer relevance of ESM1, potentially beyond.

### Limitations of the Study

In this study, we used intraportal injection to establish E0771 liver colonization, because E0771 cells poorly form spontaneous metastases when orthotopically transplanted into the mammary fat pad in an immunocompetent background. Furthermore, for the quantification of liver metastasis burden under different perturbations, we used E0771 as our sole model, as the autochthonous HCC model exhibits high variability in tumor appearance and growth. Despite this variability, the HCC model did reproduce the key histological phenotypes, supporting the generality of our findings.

## Materials and methods

### Cell culture

Mouse E0771 cells (CH3 Biosystems) were cultured in RPMI-1640 medium (ThermoFisher) supplemented with 5% FBS (Gibco), 100 IU/ml penicillin/streptomycin, 100 µg/ml amphotericin B (Lonza) and 200 nM L-Glutamine (Gibco). To trace the liver metastatic cells in vivo, E0771 cells were engineered with either YFP (addgene #111176, gift from Johannes Zuber), or firefly luciferase (addgene #19166, gift from Eric Campeau & Paul Kaufman), or H2B-HaloTag (addgene #175547, gift from Kai Johnsson).

Cells were cultured in a humidified chamber under 5% CO_2_, 37°C condition, and were tested for mycoplasma contamination regularly.

### Animal use

All animal work was done in accordance with a protocol approved by Institutional Animal Care and Use Committee (IACUC) at the Sun Yat-sen University Cancer Center (SYSUCC) (Approval Number: L025504202108022, L025504202107004 and L025504202211002).

The Cre-inducible Lyve1 overexpression (*R26^CAG-LSL-Lyve1-FLAG^*) and doxycycline-inducible Lyve1 overexpression mice (*Tigre^tetO-Lyve1-FLAG-WPRE-CAG-LSL-rtTA-WPRE^*) were generated in a C57BL/6 genetic background by inserting a tetO-Lyve1Flag-WPRE-pA-Insulator-CAG-LoxP-STOP-LoxP-rtTA-WPRE-pA expression cassette into the Igs7/TIGRE locus, respectively, using a CRISPR/Cas9-mediated genome editing system by the Shanghai Model Organisms Center (SMOC).

In brief, 5□ng□μl^−1^ Cas9 mRNA, 2□ng□μl^−1^ gRNA targeting the *Rosa26* safe harbor (5’-GGGGACACACTAAGGGAGCT -3’) or TIGRE safe harbor (5’-TAACTTTAATTCTAGCGATCTGG -3’), and 10□ng□μl^−1^ targeting vector containing 3.3□kb at both the 5’ and 3’ homologous arm of the *Rosa26* or *Tigre* target site were injected into the pronuclei of C57BL/6□J one-cell stage zygotes, which were subsequently transplanted into pseudo-pregnant female mice. The F0 chimeric offspring were then screened by PCR and were then crossed with C57BL/6□J WT mice for one generation to obtain genetically engineered F1 offspring.

C57BL/6J (WT, #000664), *Lyve1^Cre^* (#012601), Ccr2-GFP (#027619) (all from Jackson Laboratory), Tnfr1-knockout (#T013927), *Esm1^flox^* (#T016017), *Rosa26^LSL-tdTomato^* (#T002249), *H11^CAG-DreERT2^* (#T049774), *Ki67^RSR-Cre^* (#T050101) (all from _GemPharmatech), *Rosa26*_*^LSL-HA-UPRT^*_, *Rosa26*_^LSL-Lyve1-Flag^ _and *Tigre*_*^tetO-Lyve1-FLAG-WPRE-^_CAG-LSL-rtTA-WPRE_* (all from SMOC) were bred under specific pathogen free (SPF) animal facility of SYSUCC, under a 12□h-12□h light-dark cycle, at around 22□°C with around 50% humidity. The experimental group mice and littermate controls were co-housed. For genotyping, Mouse Genotyping Kit (APExBio, #K1027) was used to prepare genomic DNA. Primers used for genotyping are listed in **Supplementary Table 1**.

For retro-orbital injection of lectin, the mice were anaesthetized using an isoflurane-based small animal anesthesia unit. Otherwise, ketamine (100□mg/kg)/xylazine (10□mg/kg) were used for anesthesia. All experiments were performed at 6 to 8 weeks of age, unless otherwise indicated.

### Mouse tumor models

Mammary fat pad transplantation, portal vein injection and hydrodynamic tail vein injection were performed using the same procedures as our previous studies^28,30^.

For orthotopic breast cancer model, E0771 cells were collected from culture with 0.25% trypsin (HyClone). The cells were washed twice with PBS, counted, then resuspended in 1:1 solution of PBS and Matrigel (Phenol Red-Free and growth factor reduced; Corning). Mice were anaesthetized, and a small incision was made on the shaved abdomen to reveal the mammary gland. 5×105 E0771 cells were injected directly into the mammary fat pad. Primary breast cancer samples were harvested around day 12 to 14 post-injection.

For experimental LvMet model of breast cancer, 50 μl of E0771 cells (1×10^7^ cells/ml with PBS) were injected into the portal vein using a 31-gauge needle. Finally, the incisions of peritoneum and skin were sutured and clipped, respectively. The wound clips were removed on day 7 post-injection. LvMet samples were harvested at approximately day 21 to 24 post-injection.

For spontaneous HCC model, plasmids that overexpress Myc and knockout TP53 (Addgene #129775, #20207, and #59910) were delivered into liver by hydrodynamic tail vein injection. HCC samples were isolated at around day 30 to day 42 post-injection.

### Human tissue sample collection

Clinical samples of breast cancer LvMet were collected with informed consent from patients and were used in accordance with the approval of the Research Ethics Committee of SYSUCC (Approval Number: G2021-061-01). The samples included 15 cases of luminal B, 4 cases of HER2+, and 4 cases of TNBC, all obtained from SYSUCC. Additionally, paraffin sections of tissue arrays from HCC patient samples (N=14, #HLivH028PG01) were procured from Outdo Biotech.

### *In vivo* doxycycline and antibody treatment

For endothelial-specific Lyve1 inducible overexpression, *Lyve1^Cre^; R26^LSL-tdT^; Tigre^tetO-Lyve1-FLAG-WPRE-CAG-LSL-rtTA-WPRE^* mice were administered with 2 mg doxycycline (APExBio) by daily oral gavage, from day 14 post-portal vein injection.

For antibody-mediated blockade of Tnfr1, mice were intraperitoneally injected with 5 mg/kg anti-Tnfr1 antibody (Leinco, #T250, clone 55R-170) or the same amount of isotype-matched control antibody (Armenian hamster IgG, Leinco, #I-140, clone PIP) every other day, from day 8 after tumor engraftment, as indicated in **Fig. 4E**.

For antibody-mediated blockade of VEGF signaling, mice were intraperitoneally injected with 125 μg anti-VEGFR2 antibody (BioxCell, #BE0060, clone DC101) or the same amount of isotype-matched control antibody (Rat IgG1, BioxCell, #BE0088, clone HRPN) every other day, starting on the day of bioluminescence imaging, as indicated in **Fig. 6G**.

### Mouse tissue collection, fixation, and frozen section preparation

Whole-animal perfusion was performed. Briefly, mice were euthanized in a CO_2_ chamber, and the perfusion process started once the withdrawal reflex disappeared in pelvic limbs. Mice were perfused through left ventricle with 10 mL of cold PBS supplemented with 10 μg/mL Heparin. The perfusion was completed when the liver was decolorized. Tissues were cut into appropriate size, transferred into cold 4% PFA immediately, and were fixed overnight with gentle shaking at 4°C. After rinsing with PBS for 3 times, the tissues were immersed in PBS with 30% sucrose (w/v) at 4°C overnight. Tissues were then embedded in OCT compound (Sakura) and were immediately frozen, and 8-μm sections were cut using a Leica CM1850 cryotome.

### Scanning electron microscopy

Perfused mouse normal liver and tumor (liver metastasis and HCC) samples were promptly cut into 3-mm-diameter pieces and were fixed in 2.5% glutaraldehyde at 4°C overnight. The samples were then dehydrated in graded ethanol (50%-70%-85%-100%), acetone, and isoamyl acetate. Critical-point method was used to dry the samples (CPD300, Leica), and the dried samples were mounted onto a double-sided carbon tape. Tissues were coated with gold (MC1000 Ion Sputter Coater, Hitachi) and observed with a scanning electron microscope (Quanta 200, Thermo Fisher) at 10 kV, 5×10^−4^ Pa. Images were analyzed using FIJI software as described previously^62^. Porosity was defined as the sum of fenestrae area (μm^2^) divided by vessel area analyzed (excluding gaps, μm^2^). Fenestration frequency was defined as the number of fenestrae divided by vessel area analyzed (excluding gaps, μm^2^).

### Vascular lectin perfusion and cisplatin treatment

For the evaluation of vessel perfusion, mice were given 50 μg of DyLight 649-labeled tomato Lectin (*Lycopersicon esculentum*, Vector Labs) intravenously. After 10 minutes, mice were subjected to whole-animal perfusion, as described in “**Mouse tissue collection, fixation, and frozen section preparation**” section. Normal liver tissues, liver malignant nodules, and orthotopic breast cancer tissues were carefully removed.

For short-term cisplatin delivery assay, cisplatin (10 mg/kg, MedChemExpress, #HY-17394) was intraperitoneally injected on day 21 post-portal vein injection. After 6 hours, mice were sacrificed and perfused with PBS, and the liver tumors were collected. Harvested tissues were fixed with 4% PFA overnight at 4°C.

For long-term cisplatin chemotherapy treatment, cisplatin (2.5 mg/kg) was intraperitoneally injected one day after bioluminescence imaging and then every three days thereafter, as indicated in **Fig. 6G,H**.

### Immunofluorescence of frozen and paraffin-embedded tissue sections

Paraffin-embedded sections were deparaffinized in xylene following a graded rehydration (100%-95%-70%-50% ethanol). Antigen retrieval was performed using a steamer in heat-induced Tris-EDTA buffer (pH 9.0) for 30 min. After cooling to room temperature (RT), the slides were rinsed in water, sections were immersed in 10 mM NH_4_Cl for 5 min. Frozen sections and antigen-retrieved deparaffinized sections were rinsed in PBS, and were blocked in PBS with 0.1% Triton X-100, 1% BSA and 10% normal donkey serum for 1h at RT. Samples were incubated with following antibodies: anti-CD31 (#50408-T16, SinoBiological, 1:2000), anti-VE-cadherin (AF1002, R&D, 1:100), anti-Laminin (L9393, Sigma, 1:2000), anti-Lyve1 (AF2125, R&D, 1:100), anti-Lyve1 (14-0443-82, ThermoFisher, 1:100), anti-Ly6C (MCA2389T, BioRad, 1:100), anti-LYVE1 (AF2089, R&D, 1:100), anti-tdTomato (ARG55724, arigo, 1:500), anti-Flag (ab205606, Abcam, 1:200), anti-Esm1 (AF1999, R&D, 1:100), anti-Cisplatin DNA adducts (MABE416, Sigma, 1:200), biotinylated hyaluronic acid binding protein (#385911, Sigma, 1:250), anti-Prox1 (ab199359, Abcam, 1:500) and anti-p65 (#8242S, CST, 1:100) at 4°C overnight. Then, sections were incubated with fluorescent secondary antibodies (Alexa Fluor 488, Cy3, or Alexa Fluor 647, Jackson ImmunoResearch) that are correspondent to the species of primary antibodies at room temperature for 1h and were counterstained with DAPI (Biolegend). TrueVIEW Autofluorescence Quenching Kit (Vector Labs) was used as per manufacturer’s protocol to reduce the background. All immunofluorescence images were taken using a confocal microscope (LSM980 Zeiss).

### Tissue dissociation

To increase the yield of CD31^+^ endothelial cells, perfused mouse normal liver and tumor (LvMet and HCC) samples were dissociated into single cell suspensions using High activity normal or tumor tissue enzymatic digestion kit (RWD Life Science). Briefly, dissected tissues were rinsed with PBS, and minced into 2-mm-diameter pieces. Then, the tissues were digested in C-tubes (Miltenyi Biotech) containing RPMI-1640 medium and dissociation enzyme on a Single-cell suspension dissociator (RWD Life Science). The “Tumor_3” program was run on the dissociator, then, the tubes were incubated at 37°C with gentle shaking for 7 minutes. Above processes were repeated twice. After a final run of “Tumor_3” program, the digested suspension was passed through a 100-μm filter, mixed with Debris Removal Solution (Miltenyi Biotech) to remove debris, and 1× RBC Lysis Buffer (BioLegend) was added to lyse red blood cells, as per the manufacturer’s protocol.

### Flow cytometry

Single-cell suspensions were prepared as described in the “**Tissue dissociation**”. After lysing red blood cells, the suspension was filtered through a 70-μm filter, and washed with PBS supplemented with 1% BSA and 1.5% FBS. Cells were blocked with anti-mouse CD16/32 (1:200, clone S17011E) and Monocyte Blocker (1:200) antibodies, and stained with the following antibodies: APC anti-mouse CD31 (1:200, clone 390), APC anti-mouse CD31 (1:200, clone MEC13.3), PerCP-Cy5.5 anti-mouse CD45 (1:200, clone 30-F11), FITC anti-mouse CD4 (1:200, clone GK5.5), Pacific Blue™ anti-mouse CD8 (1:500, clone 53-6.7) (all from Biolegend), eFluor 450 anti-mouse Lyve1 (1:200, clone ALY7), PE anti-mouse Lyve1 (1:200, clone ALY7) (both from ThermoFisher). Biotinylated antibodies targeting lineage-negative markers include anti-mouse NK1.1 (clone PK136), anti-Gr1 (clone RB6-8C5), and anti-CD19 (clone 6D5) (all 1:500 Biolegend) and anti-F4/80 (CI:A3-1, BioRad, 1:500). BV421 or APC-conjugated streptavidin (1:500, Biolegend) were used to label lineage negative cells. DAPI and HelixNIR were used to exclude dead cells. Precision Count Beads (BioLegend, #424902) were used to quantify absolute cell number. Samples were analyzed using a Beckman CytoFLEX flow cytometer and data were processed using FlowJo software (v10.7).

### Cell sorting and enrichment of endothelial cells

Prior to fluorescent-activated cell sorting (FACS), CD31^+^ endothelial cells were enriched using magnetic bead-conjugated CD31 antibodies (#92-01-0143, XinBio), and subsequently passed through a magnetic sorter (XinBio), according to the manufacturer’s protocol. The enriched cells were then incubated with fluorescent antibodies (refer to “**Tissue dissociation**” and “**Flow cytometry**” sections) for FACS to purify CD45−CD31+ cells for downstream sequencing experiments.

### Bulk transcriptomic sequencing and analysis

10,000 to 20,000 DAPI−CD45−CD31+ endothelial cells were sorted using FACS. For mini-bulk RNA-seq, sequencing libraries were prepared using the Single Cell Full Length mRNA-Amplification kit (N712, Vazyme). Briefly, reverse transcription was performed to convert RNA into cDNA, followed by PCR amplification of the transcriptome, as per manufacturer’s instructions. Sequencing adapters were then added to sequence the cDNA library using the TruePrep DNA Library Prep Kit (TD501, Vazyme). The libraries were sequenced on the Illumina NovaSeq 6000 platform using 150-base pair paired-end reads.

Raw reads were trimmed using Trimmomatic (v0.39) and aligned to the indexed transcriptome (GRCm38) using STAR (v2.6.1a). Gene expression levels were quantified from the aligned reads using HTSeq (v2.0.3). The DESeq2 (v1.44.0) package in R was used to normalize reads, perform principal component analysis, and conduct differential gene expression analysis. Differential expression was performed with limma on Log2(TPM + 1) using a paired design (∼ mouseID + Tissue).

### *In situ* epigenetic library preparation

NuTRAP mice were euthanized and perfused with PBS containing 10 μg/mL Heparin (Sigma, #H3149) and 1× protease inhibitor cocktail (MCE, #HY-K0010). The nuclei were extracted with Nuclei Extraction Kit (Abbkine, #KTP4002). In brief, around 100 mg of normal liver or LvMet tissue were suspended with 1 mL of pre-chilled Lysis Buffer and mechanically homogenized. The homogenates were then incubated on ice for 20 min to facilitate nuclei release. 1 mL of pre-chilled 1.8 M Sucrose Cushion was added to each homogenate and gently mixed by pipetting. The total 2 mL of homogenate was carefully layered onto an additional 1 mL of pre-chilled 1.8M Sucrose Cushion to maintain a distinct interface. Samples were centrifuged at 4,000× *g* for 1 hour at 4°C, with acceleration and deceleration settings of 3 and 1, respectively. After centrifugation, the supernatant and intermediate debris layer was removed, and the nuclei-containing pellet was gently resuspended in 500 μL of PBS, followed by the addition of 500 μL of 2% formaldehyde in PBS. The nuclei were fixed for 5 min at room temperature and quenched by adding 2.5 M glycine to a final concentration of 125 mM, followed by an additional 5 mins incubation at room temperature with gentle agitation. Fixed nuclei were passed through a 70-μm cell strainer and centrifuged at 1,000× *g* for 10 min at 4°C. The supernatant was discarded, and the nuclei were resuspended in MACS buffer containing DAPI. 10,000 nuclei (DAPI+) expressing GFP were purified by FACS sorting for ATAC-seq and CUT&Tag-seq.

For ATAC-seq library preparation, the nuclei were resuspended in 40 μL transposase reaction mix, incubated at 37°C for 30 min on a PCR cycler, and to stop the tagmentation (NovoProtein, #N248). 160 μ L of reverse-crosslinking buffer was added to the mixture (50 μM Tris-HCl, pH7.4; 1 μM EDTA; 200 μM NaCl; 1% SDS) and incubated at 65°C overnight^63^. The samples were incubated with Proteinase K (1 μg/ml) at 55°C for 2 hours, and then purified with ChIP DNA concentrator columns (Zymo, #D5205). The DNA was eluted with 25 μL elution buffer and then amplified with PCR mix supplied with the kit.

For CUT&Tag-seq library preparation, the nuclei were resuspended in 100 μL cell membrane permeate buffer and incubated at 62°C for 10 min; 900 μL dilution buffer was then added, and samples were incubated at 37°C for 30 min (both provided in the kit #N259, NovoProtein). The ConA beads enrichment and antibody binding were performed per the manufacturer’s instructions. The reverse-crosslinking and DNA elution were similar to ATAC-seq library preparation.

The following antibodies were used for CUT&Tag assay: anti-Histone H3 tri-methyl K4 (H3K4me3, abcam, ab213224, 1:200), and goat anti-rabbit IgG secondary antibodies (abcam, #ab7085, 1:200). The PCR cycle number was optimized for each assay (ATAC-seq, 16 cycles; H3K4me3, 18 cycles). The multiplexed libraries were sequenced on Novaseq 6000 (Illumina) using 150 bp pair-end reads.

### Epigenetic sequencing data analysis

Raw reads were trimmed using Trimmomatic (v0.39) and aligned to the indexed GRCm38 genome using bowtie2 (v2.5.4). Duplicate reads and reads overlapping blacklist regions were removed using Picard (v2.19.0) and bedtools (v2.30.0), respectively. Clean BAM files were converted to bigWig files using deeptools (v3.5.3) bamCoverage using the parameters ‘--effectiveGenomeSize 2652783500 -- normalizeUsing RPKM --extendReads 200 --binSize 50 --outFileFormat bigwig’.

For principal component analysis (PCA), cleaned BAM files were used for peak calling with MACS2 (callpeak -f BAMPE -g mm -q 0.05). Reads were counted over consensus peaks with DiffBind v3.16, and differential accessibility was tested with DESeq2 using a paired sample design (∼ mouseID + Tissue) on integer-rounded counts; the condition contrast (TEC vs NEC) was extracted directly as the tested coefficient. Differentially accessible peaks were defined at FDR < 0.05 and fold change > 2. Sample relationships and quality metrics were assessed by fragment-size periodicity, fraction of reads in peaks (FRiP), TSS enrichment profiles, and PCA on variance-stabilizing-transformed (VST) counts over all consensus peaks.

Metagene heatmaps were generated in ComplexHeatmap (v2.22.0). Each analyzed region spanned 3 kb upstream of the annotated TSS (5’UTR start), the entire gene body (TSS to 3’UTR/gene end, scaled to 100 bins), and 3 kb downstream of the gene end; flanks were binned into 30 fixed 100-bp bins each (160 columns total). Minus-strand genes were flipped to 5’->3’ orientation so that the TSS aligns across rows. Read-start density per bin was extracted from each of the 16 BAM files (8 ATAC-seq, 8 H3K4me3 CUT&Tag) using restricted Rsamtools::scanBam queries, normalized to CPM, transformed to Log2(CPM + 1), and summarized per gene as mean(TEC) − mean(NEC) per bin. Rows were grouped by RNA direction: top 500 RNA-upregulated and top 500 RNA-downregulated genes (nominal *P* < 0.05, |Log2FC| > 1, ranked by *P*; genes lacking TxDb annotation were omitted, leaving 414 up / 228 down genes). Mean profiles are plotted with mean ± SEM across genes; per-bin two-sided Wilcoxon rank-sum tests between RNA-up and RNA-down groups are provided in the accompanying table.

For VEGF-axis genes and sinusoidal/capillarization markers, promoter-level heatmaps display per-sample Log2(CPM + 1) signals (z-scored) with the paired differential statistics (Log2FC, *P* value) of the strongest-count promoter-overlapping consensus peak annotated per gene. RNA-vs-epigenome concordance for the marker panel is shown as scatter plots of RNA Log2FC against promoter ATAC-seq and H3K4me3 Log2FC with linear fits. The 179 activated and 91 repressed promoter genes (Venn intersections) were additionally visualized with the whole-gene metagene heatmaps described above, with RNA Log2FC values exported in heatmap row order.

Gene sets were derived from DA peaks (gained/lost, annotated by ChIPseeker to the nearest gene and promoter-overlapping genes) and from the integrated promoter classes (activated, repressed). Over-representation analysis was performed with clusterProfiler against GO BP/MF/CC (org.Mm.eg.db; BH FDR < 0.05; background = all annotated genes) and KEGG (enrichKEGG, organism = mmu).

Genome browser tracks were visualized using WashU Epigenome Browser (https://epigenomegateway.wustl.edu/).

### Dynamic single-cell RNA sequencing and analysis of vessel isolated from mouse hepatocellular carcinoma

For dynamic scRNA-seq, 200 µl of labeling reagent containing 4-thiouridine (S^4^U) was intravenously injected into mice bearing HCC. After a 4-hour incubation period, the mice were euthanized, and whole-animal perfusion was performed as described in the section titled “**Mouse tissue collection, fixation, and frozen section preparation**”. Single-cell suspensions were prepared from freshly isolated liver or HCC tissues, following the protocol detailed in the “**Tissue dissociation**” section.

The single-cell suspensions were subjected to magnetic enrichment and subsequently stained with anti-CD45 and anti-CD31 antibodies for FACS to further purify DAPI−CD45−CD31+ endothelial cells. The sorted cells were resuspended at a concentration of 2×10^5^ cells/ml and loaded onto a microwell chip using the Singleron Matrix Single Cell Processing System (Singleron Biotech).

After barcoding the cells in the microwell, the beads were collected for the base conversion step. To achieve the base conversion of S^4^U, an iodoacetamide (IAA) alkylation reaction was performed, which introduced a carboxyamidomethyl group to the thiol group of S^4^U. This modification allows S^4^U to be recognized as cytosine during reverse transcription, resulting in T-to-C mutations in the sequencing reads. Following reverse transcription, PCR amplification, and cDNA purification, the amplified cDNA was fragmented and ligated with sequencing adapters.

Single-cell RNA-seq libraries were constructed using the GEXSCOPE Single Cell RNA Library Kit (Singleron Biotech) according to the manufacturer’s protocol. Individual libraries were diluted to 4 nM, pooled, and sequenced on the Illumina NovaSeq 6000 platform using 150 bp paired end reads.

Raw sequencing reads were processed to generate gene expression profiles using CeleScope v2.0.7 (Singleron Biotech). The CeleScope pipeline performed the following steps: (1) extraction of Barcodes and Unique Molecular Identifiers (UMIs) information from R1 reads; (2) trimming of adapter sequences and polyadenylated tails from R2 reads; (3) alignment of the reads against the GRCm38 genome assembly; (4) identification of T-to-C base conversions and differentiation between nascent and old RNA; (5) quantification of uniquely mapped reads using FeatureCounts and generate gene expression matrix.

For scRNA-seq analysis, cells with top 2% UMI counts were filtered out. The filtered count matrices were converted to Seurat (v4.3) objects. Cells expressing fewer than 200 genes or with more than 20% mitochondrial content were excluded from downstream analysis. Filtered data were then log normalized and scaled using “NormalizeData” and “ScaleData”, respectively. Cell-cell variation due to UMI counts and percent mitochondrial reads regressed out. Top 2000 variable genes were selected by “FindVariableFeatures” for principal component analysis (PCA). The top 20 principal components were used for clustering and dimensional reduction at a resolution of 1.2. The cell type identity of each cluster was determined with the expression of canonical markers based on the annotation of SynEcoSys database (https://singleron.bio/product/detail-25.html). Uniform Manifold Approximation and Projection (UMAP) algorithm was applied to visualize cells in a two-dimensional space. Violin plots displaying the expression of markers used to identify each cell type were generated by “VlnPlot”.

The “T-to-C” turn-over rate was calculated as proportion of newly synthesized transcripts in whole transcripts based on UMI counts in each gene or cell. The result was also shown by violin plot with statistical test.

### Pre-processing of clinical single cell RNA sequencing data

Three publicly available single cell RNA sequencing (scRNA-seq) datasets were obtained from previous studies of liver cancer and liver metastases (PRJCA007744)^25^, breast cancer liver metastases (SCP2702)^26^, and primary breast cancer (GSE176078)^24^.

Cell Ranger-processed gene-expression matrices aligned to the human GRCh38 reference genome and analyzed using Seurat (v4.3.1). DecontX (v1.0.0) was used to estimate ambient RNA contamination, cells with ≤100 or ≥10,000 detected genes or ≥50,000 unique molecular identifiers (UMIs) were excluded. To reduce technical variation associated with sequencing and tissue dissociation, 1,514 genes were removed, including mitochondrial (n=50), heat-shock protein (n=178), ribosomal (n=1,253), and dissociation-associated genes (n=33). scDblFinder (v1.16.0) was used to identify potential doublets. Cells with an scDblFinder score ≥ 0.8 were excluded.

Each sample was independently normalized using NormalizeData with the LogNormalize method and a scale factor of 10,000. The top 3,000 highly variable genes were identified for each sample using FindVariableFeatures with the variance-stabilizing transformation (vst) method. Genes were ranked by the frequency with which they were identified as highly variable across samples, and the top 2,000 recurrently variable genes were selected.

During data integration, batch effects across samples and datasets were corrected using Harmony (v1.2.0), with sample identity specified as the batch variable and a maximum of 10 iterations. A shared nearest-neighbor graph was constructed using the first 30 Harmony dimensions, followed by Louvain clustering at resolutions ranging from 0.4 to 1.6 in increments of 0.2. UMAP based on the same Harmony dimensions was used for visualization.

### Cell-type annotation and endothelial cell subclustering

Major cell lineages were annotated according to canonical marker-gene expression. T and NK cells were identified using CD3D, CD3E, CD4, CD8A, FOXP3, TRDC, NKG7, and KLRD1; B cells using CD79A and MS4A1; monocytes and macrophages using CD14, FCGR3A, CD68, and CD163; dendritic cells using CD1C and LAMP3; mast cells using TPSAB1; neutrophils using CSF3R and S100A8; endothelial cells using VWF and PECAM1; fibroblasts using COL1A1 and COL1A2; and epithelial cells using EPCAM and tissue-specific epithelial markers such as ALB.

For detailed subtype annotation, cells belonging to each major lineage were extracted from the integrated dataset and analyzed separately. Within each lineage, normalization, highly variable gene selection, data scaling, PCA, Harmony batch correction, neighbor-graph construction, clustering, and UMAP visualization were repeated using the same general workflow described above. Cell subtypes were assigned according to established lineage-specific marker genes. Clusters lacking canonical lineage markers or showing conflicting markers from multiple lineages were further evaluated as potential low-quality cells, contaminants, or doublets and excluded where appropriate.

### RNA velocity analysis

Spliced and unspliced transcript counts were quantified from Cell Ranger-generated BAM files using Velocyto (v0.17.17) with the GRCh38 gene annotation and corresponding repeat-mask file. Only barcodes retained in the filtered feature-barcode matrices were included.

RNA velocity analysis was performed using scVelo (v0.3.3). Endothelial cells were selected according to their cell barcodes, and previously generated cell-type annotations and UMAP coordinates were transferred to the AnnData object. Genes with fewer than 50 shared counts were filtered out, and the top 500 highly variable genes were retained. Expression moments were calculated using 50 principal components and 50 nearest neighbors. Gene-specific transcriptional dynamics were fitted using scvelo.tl.recover_dynamics, and RNA velocities were estimated in dynamical mode using scvelo.tl.velocity. The velocity graph was constructed based on cosine similarities using scvelo.tl.velocity_graph. The inferred velocities were projected onto the precomputed UMAP embedding and visualized as streamlines. Latent time was calculated using scvelo.tl.recover_latent_time and visualized on the same UMAP embedding.

### Pseudotime trajectory analysis

Endothelial cell trajectories were reconstructed using Monocle3 (v1.3.1). Raw RNA counts and cell metadata were extracted from the Seurat object and used to construct a cell_data_set. Data were preprocessed by PCA using 20 dimensions, and the Harmony-corrected UMAP coordinates generated in Seurat were imported to ensure consistency between analyses. Cells were clustered using cluster_cells, and the principal trajectory graph was learned using learn_graph. LSEC cells were defined as the root state, and cells were ordered along pseudotime using order_cells. Trajectories, pseudotime distributions, and gene-expression dynamics were visualized using plot_cells and plot_genes_in_pseudotime.

### Spatial transcriptomics processing and regional analysis

Spatial transcriptomic data of colorectal cancer liver metastasis and hepatocellular carcinoma were obtained from OEP00175664 and HRA000437^65^, respectively. Corresponding histological images were reviewed by pathologists, and samples containing both normal liver tissue and liver metastases or primary liver tumors were selected. Based on histopathological features, spatial spots were classified into normal, tumor-border, and tumor regions.

Space Ranger (v3.1.2) was used to convert Visium spatial transcriptomics raw counts into feature-barcode HDF5 matrices. Filtered matrices and the corresponding spatial information were loaded into Seurat using Load10X_Spatial, with gene-expression data stored in the Spatial assay. Histopathology-based regional annotations were assigned to individual spots according to their barcodes, and their spatial distributions were visually inspected using SpatialDimPlot. Sample identity was retained in the metadata for subsequent comparisons.

Six spatial transcriptomics datasets were merged into a single Seurat object and normalized using SCTransform (v0.4.3). Principal component analysis was subsequently performed, and a shared nearest-neighbor graph was constructed using the first 30 principal components. Spatial spots were clustered using the Louvain algorithm implemented in FindClusters at a resolution of 0.5. UMAP was generated using the same 30 principal components to visualize the transcriptional relationships among spots. The quality control analyses of these six datasets are provided in **Supplementary Table 2**.

To assess regional variation in inflammatory gene expression, normalized expression values were extracted for each spatial spot. Spots were grouped by sample and histopathological region, and the mean expressions of selected inflammatory mediators were calculated separately for normal, border, and tumor regions. Regional expression patterns were visualized for each sample using line plots.

Cell-type composition was estimated using cell2location (v0.1.4). An annotated scRNA-seq dataset was used as the reference, and genes were selected using filter_genes with cell_count_cutoff = 60, cell_percentage_cutoff2 = 0.00015, and nonz_mean_cutoff = 1.50. The reference model was trained for 300 epochs, and posterior distributions were estimated using 1,000 samples. The expected number of cells per spot was set to 30, and detection_alpha was set to 200 for cell2location model fitting. The model was trained for 30,000 epochs, and posterior cell-type abundances were estimated using 1,000 samples. The 5% posterior quantile was used for spatial visualization.

### TCGA Pan-cancer analysis of ESM1

The processed scRNA-seq data integrated from 31 cancer types were obtained from PRJCA018695. The processed Seurat objects were downloaded from the online data browser (http://resource.yin-lab.com/Panvascular/). Pseudotime expression analysis of ESM1, VEGFA, and ANGPT2 was performed using the built-in RStudio Shiny tool in that online data browser.

The ESM1 expression level (‘rna_seq_v2_mrna_median_all_sample_Zscores’) and clinical annotations from TCGA dataset were downloaded from cBioportal using ‘getDataByGenes’ and ‘clinicalData’ functions in “cBioPortalData” R package. Hazard ratios were calculated using the Cox proportional hazards model.

### Statistical analysis

Sample sizes are denoted in figures or figure legends. Data were presented as mean ± SEM, if not otherwise specified. For the dot plots of immunostaining results, smaller dots without an outline are values from individual fields, and circles with an outline represent mean values taken over 3−5 fields from the same mouse, unless otherwise noted. The statistical tests were performed by comparing the individual animals. All statistical analyses were performed with Prism 9 software (GraphPad) or R (version 4.3.1). Two-tailed Student’s *t* test or Mann-Whitney *U*-test was used to compare two groups. Statistical tests were denoted in the figure legends. *P* values lower than 0.05 were considered statistically significant.

## Data availability

Materials and additional details will be available upon request, contingent upon material transfer agreements with SYSUCC. The raw sequencing data and processed data matrices in this study have been deposited at the China National Center for Bioinformation (PRJCA075202) and will be accessible prior to publication.

## Code availability

Custom codes used in this study have been deposited at GitHub and are available at https://github.com/chuangzhao0601/Liver-Metastasis-Vascular-Normalization.git.

## Acknowledgement

L.T. is supported by the General Program of the National Natural Science Foundation of China (NSFC) (Grant No. 82573221), the Foundation for Innovative Research Groups of the NSFC (Grant No. 82521103), and the Young Talents Program of SYSUCC (Grant No. YTP-SYSUCC-0045).

## Contributions

H-Y.H. and L.T. conceived the idea and performed most of the experiments. C.Z. and L.T. performed bioinformatic analyses. X-N.Z. performed NuTRAP in situ epigenetic profiling. X-N.Z. and J-M.Z. assisted with immunofluorescence staining and quantification. Y-X.S. collected liver metastasis tissue samples from breast cancer patients. L.T. wrote the manuscript. All co-authors have read and approved the manuscript.

