## Supplementary Fig. 1-6 for "Targeting the sinusoidal-to-capillarized transition to normalize vessels in breast cancer liver metastasis"

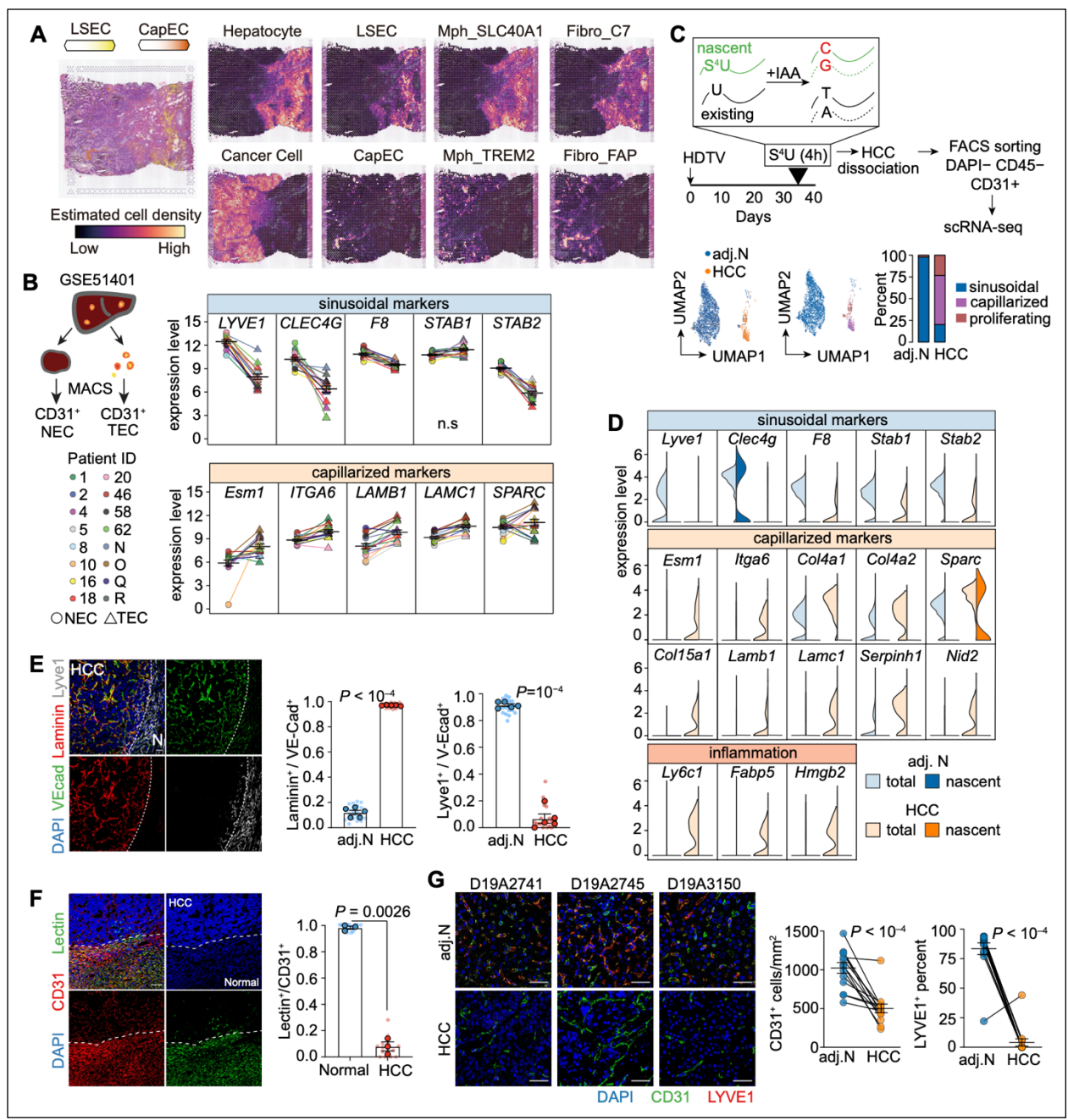

**Supplementary Figure S1. Loss of sinusoidal and gain of capillarized features in vessels of other liver malignancies**

**A.** Spatial transcriptomic analysis of colorectal cancer liver metastasis samples. Expression of cell-type markers was overlaid onto H&E staining. Sinusoidal gene expression was enriched in hepatocyte-containing normal liver regions, whereas capillarized endothelial cells appeared in the carcinoma region.

- **B.** Alterations in sinusoidal and capillarized genes in magnetic-activated cell sorting (MACS)-enriched CD31<sup>+</sup> tumor endothelial cells (TECs) and CD31<sup>+</sup> normal endothelial cells (NECs) from HCC patients.
- **C.** (Top) Schematic of dynamic single-cell RNA sequencing (scRNA-seq). Nascent RNA labeled with 4-thiouridine (S4U) was distinguished from mature RNA through iodoacetamide (IAA)-mediated alkylation and subsequent bioinformatic analysis. (Bottom) Uniform manifold approximation and projection (UMAP) embedding of endothelial cells from HCC and adjacent normal (adj. N) tissue, with stacked bar plots showing the composition of endothelial subsets.
- **D.** Violin plots showing expression levels of sinusoidal, capillarized, and inflammatory genes in nascent RNA and total RNA from HCC and adj. N liver tissues.
- **E.** Immunostaining showing vascular laminin deposition and Lyve1 expression in HCC samples and adj. N liver tissues (scale bar, 50 μm).
- **F.** Lectin perfusion assay showing dysfunctional perfusion of TECs in HCC (scale bar, 50 μm).
- **G.** Immunostaining of clinical HCC samples showing loss of the sinusoidal marker Lyve1 in TECs (scale bar, 50 μm).

Mean ± s.e.m. shown. Smaller dots represent values from individual fields, and outlined circles indicate mean values taken over multiple fields/sections from the same mouse (**E,F**). *P* values were calculated using two-tailed unpaired Mann-Whitney *U*-test (**D**), and paired Student's *t*-test (**B, E-G**).

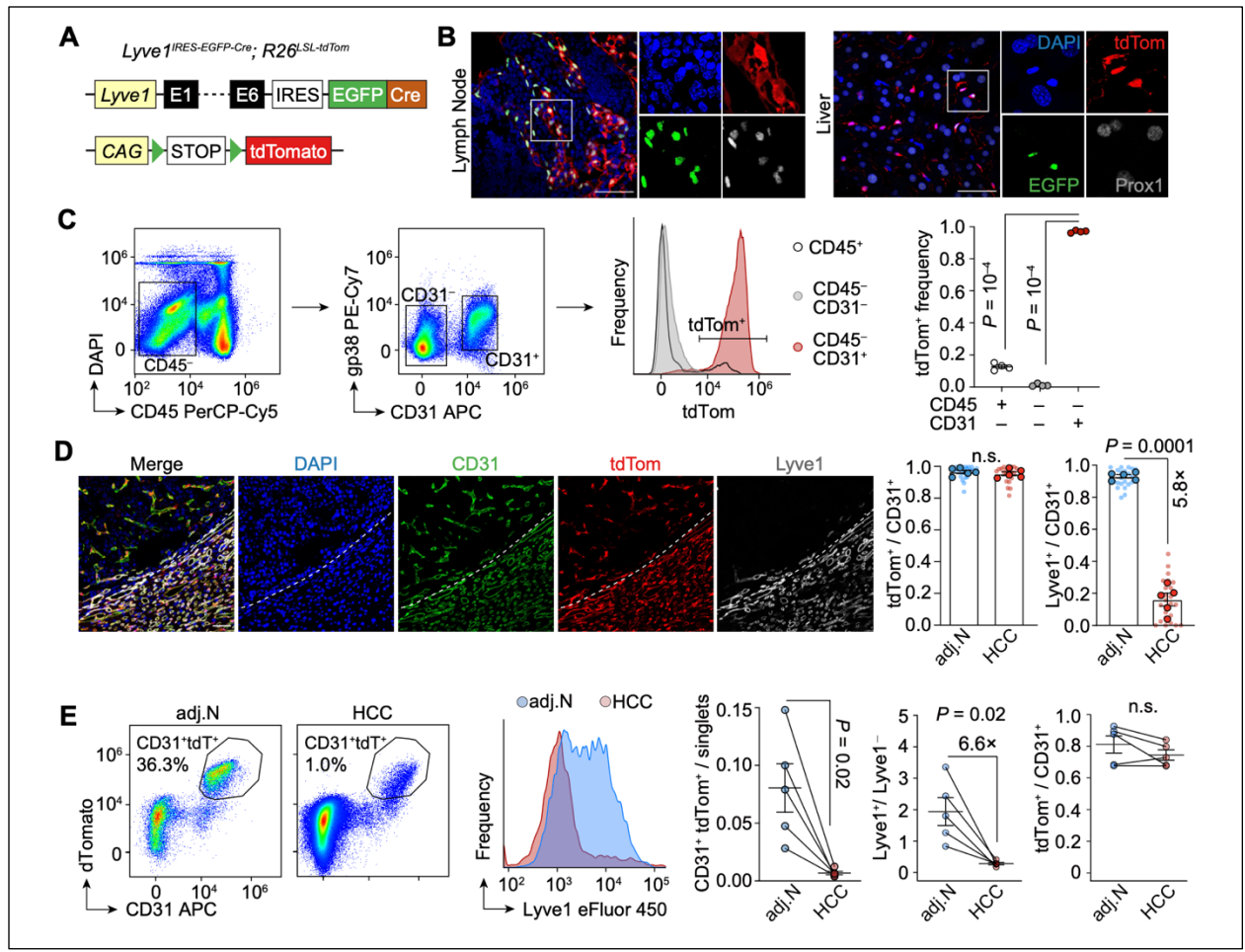

### Supplementary Figure S2. Efficiency and specificity of the sinusoidal lineage tracer and its application in HCC

**A.** Schematic of LSEC tracing using *Lyve1*Cre. When crossing with a red fluorescent protein (tdTomato) reporter (*R26<sup>LSL-tdTomato</sup>*), CAG-promoter-driven tdTomato expression is activated, enabling tracing of LSEC descendants even after *Lyve1* expression is repressed.

**B.** Immunostaining of the lymphatic vessel marker Prox1 showing that tdTomato+ cells in the liver of *Lyve1<sup>Cre</sup>; R26<sup>LSL-tdTomato</sup>* mice are not lymphatic endothelial cells, confirming the specificity of this model for tracing LSECs (scale bar, 50 μm).

**C.** Flow cytometry analysis of tdTomato+ cells. More than 95% of DAPI- CD45- CD31+ endothelial cells were tdTomato+, demonstrating the efficiency of the model.

**D,E.** *Lyve1*-lineage tracing of the autochthonous mouse hepatocellular carcinoma model, phenocopying the results observed in the breast cancer LvMet model. Immunostaining (**D**) and flow cytometry (**E**) show that the percentage of tdTomato+ cells among CD31+ endothelial cells is unchanged despite significantly decreased *Lyve1* expression.

Mean ± s.e.m. shown. Smaller dots represent values from individual fields, and outlined circles indicate mean values taken over multiple fields/sections from the same mouse (**D**). *P* values were calculated using two-tailed paired Student's *t*-test (**C-E**).

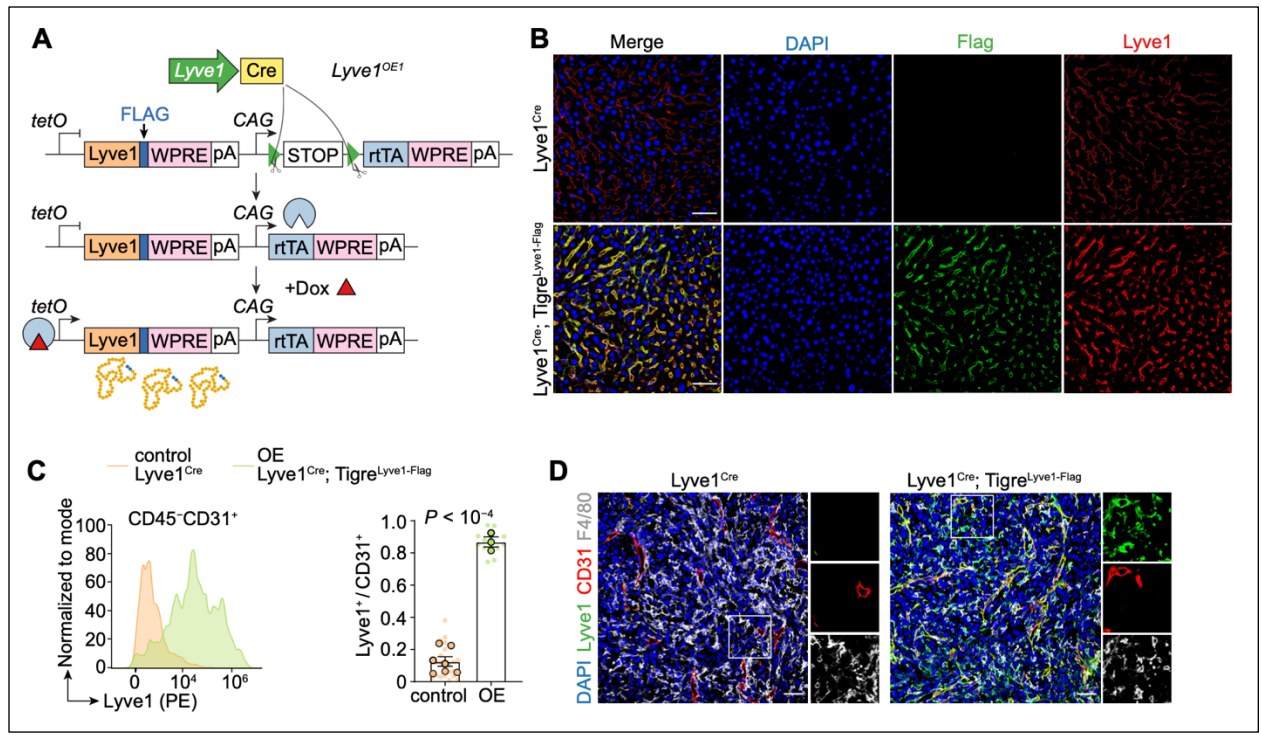

#### Supplementary Figure S3. A doxycycline-inducible Lyve1 overexpression model exhibits leaky expression in LvMet

**A.** Schematic of the endothelial-specific conditional Lyve1 overexpression (*Lyve1*<sup>OE1</sup>) mice (genotype: *Lyve1*<sup>Cre</sup>; *R26*<sup>LSL-tdT</sup>; *Tigre*<sup>tetO-Lyve1-Flag-WPRE-CAG-LSL-rtTA-WPRE</sup>).

**B,C.** Immunostaining (**B**) and flow cytometry (**C**) of Lyve1 in *Lyve1*<sup>OE1</sup> mice and littermate controls lacking the TIGRE transgenic allele (scale bar, 50 μm). Flag expression indicates ectopic Lyve1 expression, which can be distinguished from endogenous untagged Lyve1.

**D.** Immunostaining showing non-endothelial expression of Flag-tagged Lyve1 in F4/80<sup>+</sup> macrophages, suggesting Dox-independent, nonspecific ectopic Lyve1 expression in *Lyve1*<sup>OE1</sup> mice.

Mean ± s.e.m. shown. Smaller dots represent values from individual fields, and outlined circles indicate mean values taken over multiple fields/sections from the same mouse (**C**). $P$ value was calculated using two-tailed unpaired Student's $t$-test (**C**).

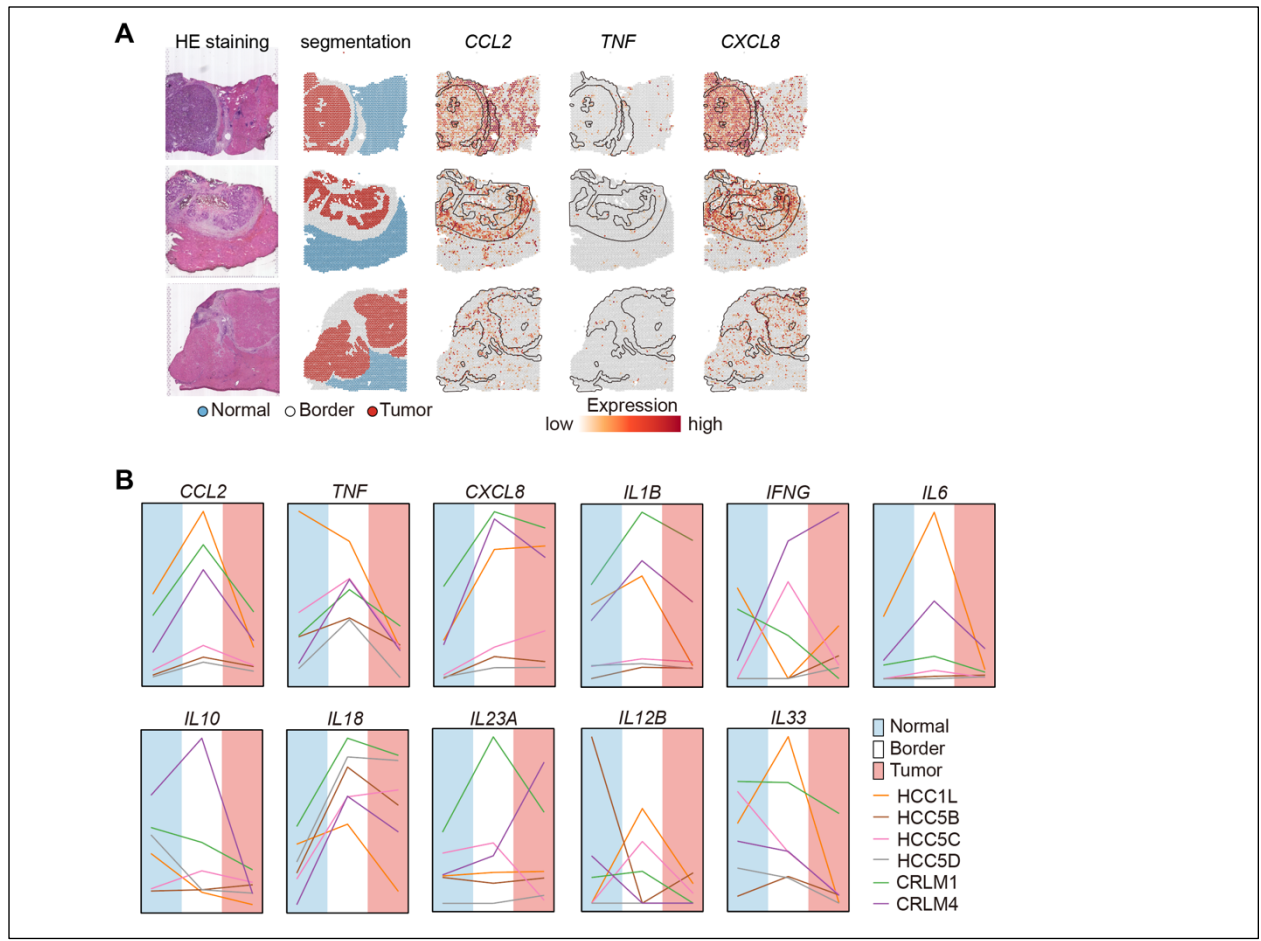

#### Supplementary Figure S4. Spatial transcriptomic analysis of inflammatory factors in liver malignancies

**A.** Representative images showing H&E staining, computational segmentation of normal liver, carcinoma, and border regions, and color-coded expression of CCL2, TNF, and CXCL8.

**B.** Line plots showing the changes in 11 inflammatory factors (cytokines and chemokines) across six datasets, including hepatocellular carcinoma (HCC) and colorectal cancer liver metastasis (CRLM).

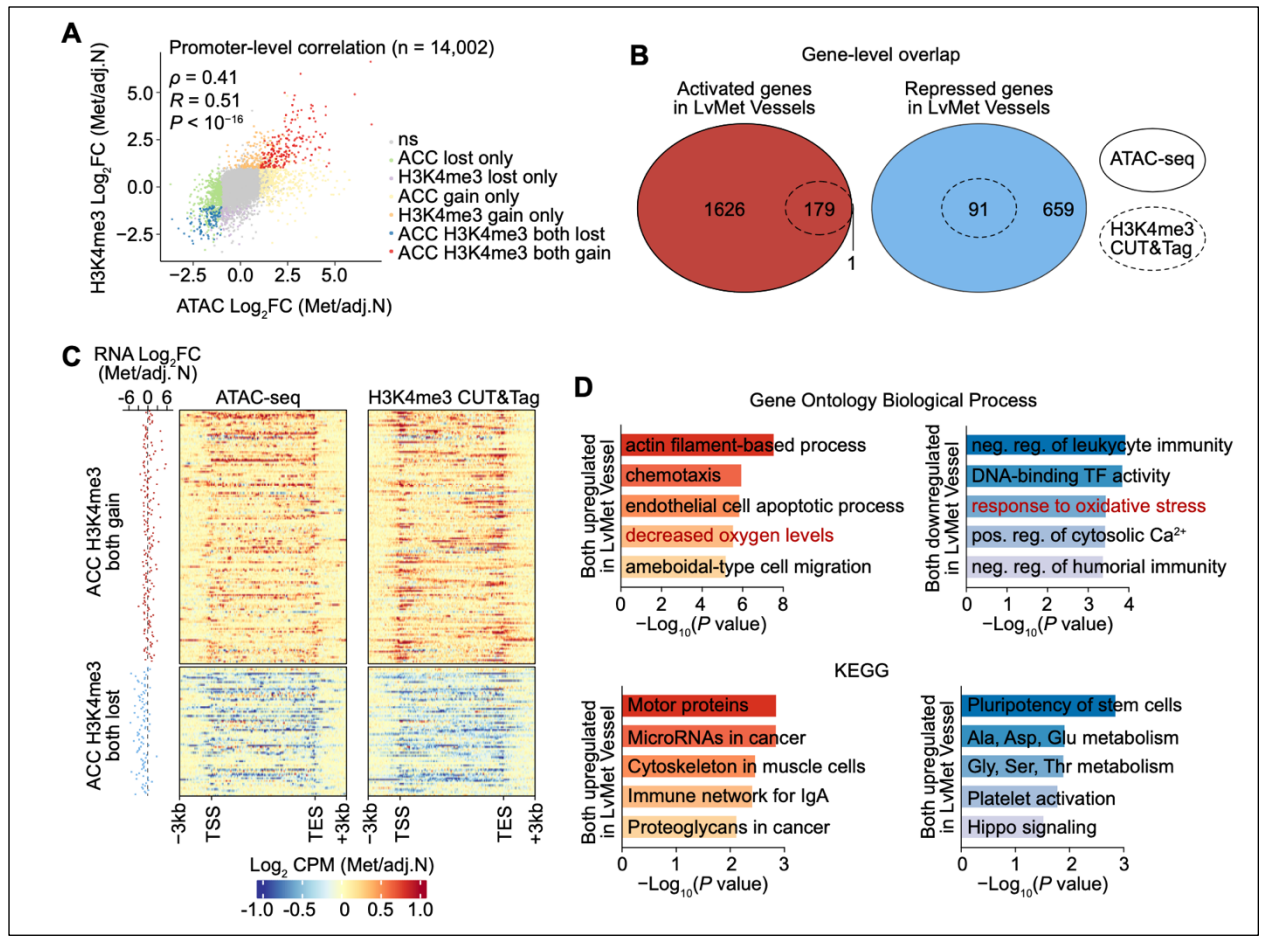

#### Supplementary Figure S5. Integrated analysis of ATAC-seq and H3K4me3 CUT&Tag

**A.** Quadrant scatter plot of promoter accessibility Log<sub>2</sub>FC (x-axis) versus promoter H3K4me3 Log<sub>2</sub>FC (y-axis) for the 14,002 promoters. Each dot is one promoter, colored by different category.

**B.** Overlap of gene-level accessibility changes and H3K4me3 changes for activated (left) and repressed (right) genes. H3K4me3 changes are almost entirely nested within accessibility changes, indicating that gained accessibility is necessary but not sufficient for H3K4me3 promoter gain.

**C.** Promoter signal (Z-scored Log<sub>2</sub> CPM) heatmap for ATAC-seq (left) and H3K4me3 (right) across the 179 activated (top) and 91 repressed (bottom) promoter genes identified in **(B)**. The RNA Log<sub>2</sub>FC (LvMet vs adj.N) of each gene is shown as the dot plot at left.

**D.** KEGG and Gene Ontology analysis of the activated (n = 179) and repressed (n = 91) promoter gene sets.

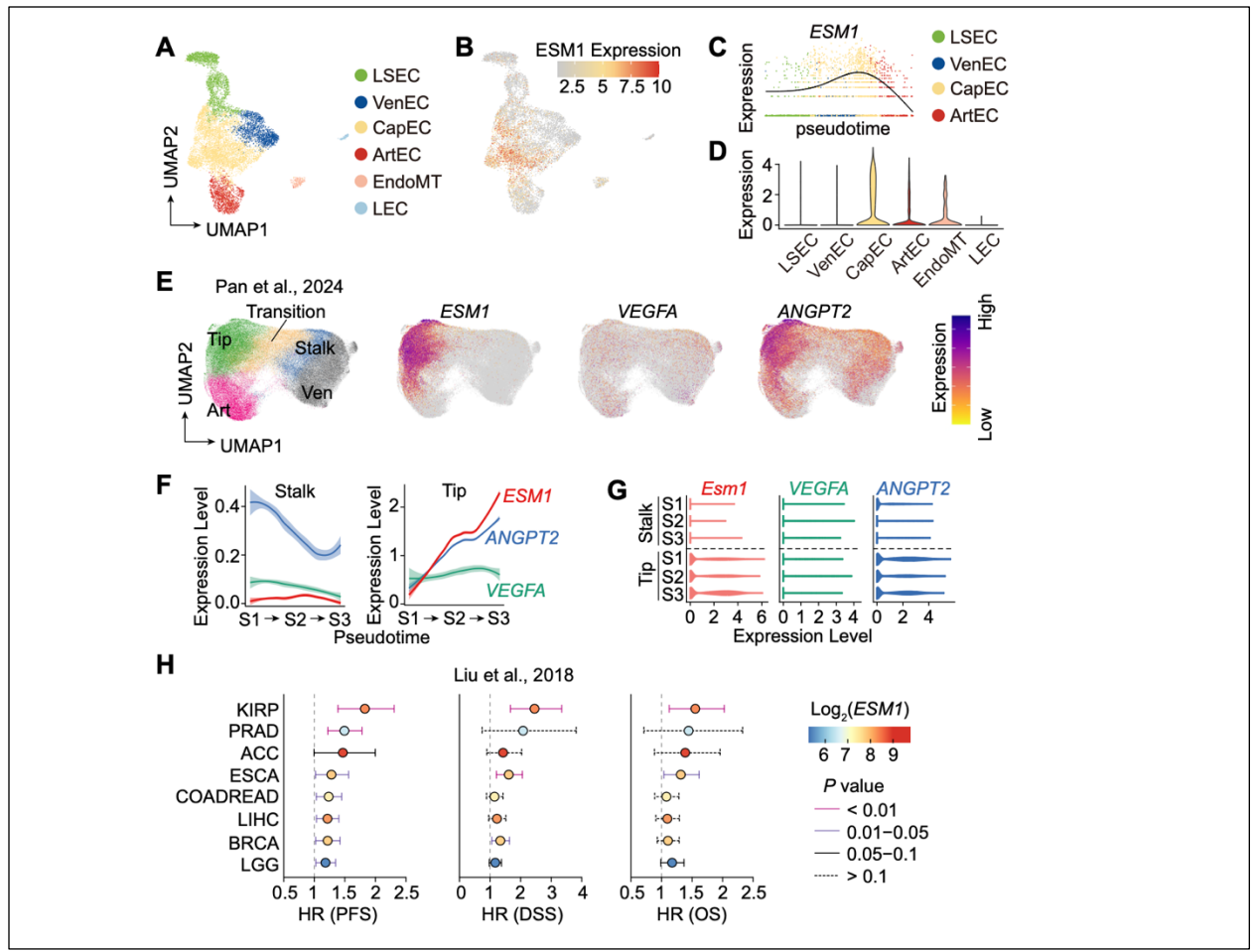

#### Supplementary Figure S6. Clinical relevance of ESM1 across multiple cancer types

**A–D. Expression of ESM1 in the integrated endothelial scRNA-seq dataset** comprising normal liver, hepatocellular carcinoma, liver metastasis of breast cancer, and primary breast cancer. The UMAP colored by endothelial subset (**A**) is shown as a reference. The feature plot (**B**), pseudotime curve (**C**), and boxplot (**D**) show that *ESM1* expression peaks in capillarized endothelial cells.

**E.** Feature plots from scRNA-seq of tumor vessels showing the expression levels of *ESM1*, *VEGFA*, and *ANGPT2* across TEC subsets.

**F,G.** Pseudotime analysis (**F**) and violin plots (**G**) showing the expression patterns of *ESM1*, *VEGFA* and *ANGPT2* along the differentiation trajectories of stalk cells and tip cells.

**H.** Survival analysis of ESM1 expression across multiple cancer types. The color indicates the mean ESM1 expression level in each cancer type. *P* values were calculated using Cox proportional hazards models, and statistical significance is represented by the color or line type of the error bars (PFS, progression-free survival; DSS, disease-specific survival; OS, overall survival; KIRP, kidney renal papillary cell carcinoma; PRAD, prostate adenocarcinoma; ACC, adrenocortical carcinoma; ESCA, esophageal carcinoma; COADREAD, colon adenocarcinoma/rectum adenocarcinoma; LIHC, liver hepatocellular carcinoma; BRCA, breast invasive carcinoma; LGG, brain lower grade glioma).
