## Supplementary Table 1-2 for "Targeting the sinusoidal-to-capillarized transition to normalize vessels in breast cancer liver metastasis"

| Strain | PrimerName | Sequence (5'->3') | Band Size |
| --- | --- | --- | --- |
| Lyve1-Cre<br>JX #012601 | WT_F | TGCCACCTGAAGTCTCTCCT | WT: 425 bp<br>KI: 210 bp |
|  | KI_F | GAGGATGGGGACTGAAACTG |  |
|  | Com_R | TGAGCCACAGAAGGGTTAGG |  |
| R26-LSL-tdT<br>GemPharm #T002249 | WT_F | CCCAAAGTCGCTCTGAGTTGTTA | WT: 479 bp<br>KI: 317 bp |
|  | WT_R | TCGGGTGAGCATGTCTTTAATCT |  |
|  | KI_F | CGGCATGGACGAGCTGTACAAG |  |
|  | KI_R | TCAGCAAACACAGTGCACACCAC |  |
| H11CAG-DreERT2<br>GemPharm #T049774 | Com_F | CAGCAAAACCTGGCTGTGGATC | WT: 412 bp<br>KI: 304 bp |
|  | WT_R | ATGAGCCACCATGTGGGTGTC |  |
|  | KI_R | CCTCCTCTCCTGACTACTCCCAGTC |  |
| Ki67RSR-Cre<br>GemPharm #T050101 | Com_F | CAGAGCTAACTTGCGCTGACTGGA | WT: 341 bp<br>KI: 649 bp |
|  | WT_R | CCCGATTCCATTTGGAAGCTC |  |
|  | KI_R | TAGAAGGCACAGTCGATCCTCTAG |  |
| Rosa26LSL-HA-UPRT<br>custom generated | Com_F | TCAGATTCTTTTATAGGGGACACA | WT: 967 bp<br>KI: 1215 bp |
|  | WT_R | TAAAGGCCACTCAATGCTCACTAA |  |
|  | KI_R | AAGTAGCAAAAGAGGAAAGAGAGACA |  |
| Lyve1-OE1 (Dox-inducible)<br>custom generated | WT_F | GCTTCCCCAACGGTCACTTA | WT: 461 bp<br>KI: 237 bp |
|  | WT_R | TCCCAAGTCATCCCTCCCAT |  |
|  | KI_F | CCTGGAGAAACAGCTAAAG |  |
|  | KI_R | CATAGCGTAAAAGGAGCAACA |  |
| Lyve1-OE2 (R26-LSL-Lyve1) | WT_F | TCAGATTCTTTTATAGGGGACACA | WT: 967 bp<br>KI: 282 bp |
|  | WT_R | TAAAGGCCACTCAATGCTCACTAA |  |
|  | KI_F | ACCCAAAACTACGGTGCGAT |  |
|  | KI_R | GGCCACAACTCCTCATAAAG |  |
| Ccr2 GFP<br>JX #027619 | Com_F | AATAATCATTTTGTTCTCTGACCAC | WT: 514 bp<br>KI: 150 bp |
|  | WT_R | ACAGCATGAACAATAGCCAAGT |  |
|  | KI_R | CTGAAC TTGTGGCCGTTTAC |  |
| Tnfr1 KO<br>GemPharm #T013927 | WT_F | CTCCCTTCAACTTGATTTCCC | WT: 451 bp<br>KI: 288 bp |
|  | WT_R | CTTGGTCTCTGGTGGCTCTGT |  |
|  | KI_F | CTTCCTTTTGTTTCTGTCCCG |  |
|  | KI_R | CTCTAAAGCAGCAACGCCAAC |  |
| Esm1 flox<br>GemPharm #T016017 | PCR_F | AGAAGCATTTGAAGTGTGCTGATCC | WT: 492 bp<br>KI: 593 bp |
|  | PCR_R | AACGTTATTGCTTGCACTGCTGAG |  |
|  | Seq_F | GTTTCACGATATTCCCTTAGGGAGTG |  |

| Regional spot counts and QC metrics of Visium samples |  |  |  |  |  |  |  |
| --- | --- | --- | --- | --- | --- | --- | --- |
| Study | Sample ID | Region | Number of spots | Fraction of analyzed spots | Median genes per spot | Median UMI counts per spot | Total Genes Detected |
| PMID:<br>34417225<br>Ref #64 | lvmet1 | Normal | 1647 | 0.4304757 | 4694 | 16963 | 23029 |
|  |  | Border | 664 | 0.1735494 | 5419 | 19801 | 21499 |
|  |  | Tumor | 1515 | 0.3959749 | 6455 | 29506 | 23189 |
|  | lvmet4 | Normal | 1601 | 0.4302607 | 982 | 1844 | 19412 |
|  |  | Border | 1161 | 0.3120129 | 2217 | 4573 | 20370 |
|  |  | Tumor | 959 | 0.2577264 | 2381 | 5104 | 19996 |
| PMID:<br>34919432<br>Ref #65 | HCC-1L | Normal | 1261 | 0.4518094 | 3073 | 10803 | 20010 |
|  |  | Border | 447 | 0.1601576 | 3518 | 10684 | 19071 |
|  |  | Tumor | 1083 | 0.388033 | 6181 | 33525 | 21443 |
|  | HCC-5B | Normal | 422 | 0.1066195 | 1086 | 2169 | 15624 |
|  |  | Border | 1032 | 0.2607377 | 1495.5 | 2811.5 | 18162 |
|  |  | Tumor | 2504 | 0.6326427 | 1972 | 4411 | 19902 |
|  | HCC-5C | Normal | 768 | 0.203336 | 2289 | 7666 | 17888 |
|  |  | Border | 886 | 0.2345777 | 1567.5 | 3108.5 | 18195 |
|  |  | Tumor | 2123 | 0.5620863 | 2084 | 5720 | 19648 |
|  | HCC-5D | Normal | 721 | 0.165671 | 1698 | 4694 | 17165 |
|  |  | Border | 1031 | 0.2369026 | 1593 | 3660 | 18253 |
|  |  | Tumor | 2600 | 0.5974265 | 2052.5 | 5849.5 | 19989 |
